# Evidence of Adaptation in Structural Variants among Wild Populations of the purple sea urchin, *Strongylocentrotus purpuratus*

**DOI:** 10.64898/2026.01.20.700628

**Authors:** Csenge Petak, Daniel E. Sadler, Melissa H. Pespeni, Joaquin C. B. Nunez

## Abstract

Structural variants (SVs) are increasingly recognized as important components of genetic architecture, complementing and extending beyond traditionally studied single nucleotide polymorphisms. Though growing, our understanding of the evolutionary forces maintaining SVs in natural populations is limited. Chromosomal inversions in particular can facilitate local adaptation in populations with high gene flow, including many marine species. The purple sea urchin (*Strongylocentrotus purpuratus*) is a powerful system to study these dynamics due to its high gene flow, lack of population structure, and broad latitudinal range. We analyzed whole genome sequence data from 137 individuals from seven populations to identify structural variants using local PCA, linkage disequilibrium, and *F*_ST_ analyses. We integrated Bayesian selection scans and population genetic statistics to test for signatures of selection in putative inversions. We identified nine loci showing signatures consistent with inversion polymorphisms, including three-way genotype clustering, long range linkage, and the characteristic hanging-bridge pattern. These loci are polymorphic within sites and along the species range with three loci showing concordant signatures of selection based on enrichment of *X*^*T*^*X* outliers and distinct patterns of allelic age consistent with positive and balancing selection. In addition, these loci show enrichment for genes associated with biomineralization and development. Our results are the first instance of identifying putative inversions in the purple sea urchin, adding to the genomic repertoire of this model species. Our results add to the growing evidence that chromosomal inversions are a key component of standing genetic variation in natural populations with an important role in adaptation to heterogeneous environments.

**Significance statement:** Chromosomal inversion polymorphisms are an important part of the repertoire of standing genetic variation in wild populations and can facilitate adaptation in the face of strong gene flow. The purple sea urchin, a widely studied marine species, exhibits high gene flow, large population sizes, and extensive genetic diversity across diverse environmental conditions, making it an ideal model for evolutionary genomics. We identified nine putative inversions, three with signatures of selection, adding echinoderms to the growing list of phyla with putatively adaptive inversions. These findings provide new insights into structural variation in a highly dispersive marine species and highlight potential evolutionary mechanisms maintaining these polymorphisms.

## INTRODUCTION

A central challenge in evolutionary biology is understanding the interplay between gene flow, which acts to homogenize populations, and natural selection, which can maintain adaptive differentiation (Maruki et al., 2022; Slatkin, 1987; Yeaman and Whitlock, 2011). Indeed, in classical population genetics theory, gene flow and selection can be viewed as antagonistic forces. Under this framework, adaptive evolution may be inhibited when migration overwhelms natural selection (Felsenstein, 1976), specifically when selection is too weak to preserve advantageous alleles attempting to become spatially structured (Haldane, 1930, *but see* Kottler et al., 2021). In this context, the balance between gene flow and selection is often described through the forces of gene swamping and migration load (Bolnick and Nosil, 2007; Lenormand, 2002). Two processes that stem from the movement of maladapted alleles in structured metapopulations, which can obscure signals of adaptation across complex spatial scales, particularly for traits with polygenic architectures (Yeaman, 2015).

Despite these theoretical expectations, empirical evidence in the era of genomics have highlighted that adaptive differentiation can be quantified even among populations connected by high levels of gene flow (i.e., weakly structured; Comeault et al., 2015; Joron et al., 2011; Laurent et al., 2016; Pespeni et al., 2012, 2010). Across these studies, chromosomal inversions are a prevalent form of genetic variation that can facilitate the maintenance of adaptative divergence even in the face of high gene flow (Joron et al., 2011; Kirubakaran et al., 2016; Nunez et al., 2024; Twyford and Friedman, 2015). For example, adaptive inversions can contribute to the maintenance of local adaptation by protecting beneficial mutations from gene swamping (Schaal et al., 2022). Furthermore, through the suppression of recombination between alternative karyotypic classes, inversions can maintain sets of co-adapted alleles in strong linkage, ensuring they will be inherited together and preventing the fitness loss that would otherwise arise in the offspring of locally adapted parents (Charlesworth, 2016; Dobzhansky and Wright, 1943; Kapun et al., 2023; Kapun and Flatt, 2019; Yeaman et al., 2016).

Natural populations of the purple sea urchin, *Strongylocentrotus purpuratus*, are a robust system to assess the role of chromosomal inversions in the interplay of natural selection and high gene flow. *S. purpuratus* is an inter- and sub-tidal marine invertebrate that lives in coastal rocky reefs along a broad latitudinal gradient from the cold waters of Alaska to the warmer waters of Baja California, Mexico, serving as a natural laboratory to study adaptation across heterogeneous environments. *S. purpuratus* are long-lived (> 50 years, Amir et al., 2020), dioecious broadcast spawners, releasing their eggs and sperm into the water column where fertilization occurs. Resulting feeding planktonic larvae can spend weeks to months traveling the coastal currents before recruiting to a suitable place to live. This high dispersal potential can swamp local adaptation but also suggests that selection on dispersing larvae is an important mechanism for limiting phenotype-environment mismatching (Levene, 1953; Marshall et al., 2010; Pespeni et al., 2013). As adults, *S. purpuratus* also has a disproportionate impact on its habitat as a voracious feeder on macroalgae. Indeed, their overgrazing of biodiverse kelp forests can result in the formation of desolate urchin barrens (Filbee-Dexter and Scheibling, 2014). *S. purpuratus* has also been a model organism in developmental biology for over a century resulting in a high-quality, well-annotated genome (Sodergren et al., 2006). Previous population genomic studies along the west coast of North America have revealed high levels of genetic diversity, gene flow, and a lack of population structure across its distribution, making them a robust model to test for signatures of natural selection along the species’ range (Pespeni et al., 2010, 2012, 2013; Petak et al., 2023). Moreover, experimental selection and physiology studies have revealed ample functional phenotypic and genetic variation in the urchin genome, the raw material for spatially heterogeneous selection (Hedrick, 2006; Nunez et al., 2020; Pespeni et al., 2013b, 2013a). These patterns of adaptive differentiation, in the context of the species’ high levels of gene flow, raise the question of whether urchin genomes harbor chromosomal inversions and whether such structural variants contribute to the patterns of local adaptation reported in this species.

In this study, we conducted the first genome-wide scan for structural variants including potentially adaptive inversions in natural populations of *S. purpuratus*. We leveraged an extensive panel of genomic variation sampled across a broad portion of the urchin’s habitat, consisting of 137 genomes from single individuals. Specifically, we applied a local principal component analysis approach to identify putative inverted regions across the genome and to pinpoint inversion-informative markers associated with these structural variants (Ma and Amos, 2012, Li and Ralph 2019). Using these data, we aimed to (1) identify patterns of genomic variation consistent with structural variants or putative inversions associated with reduced recombination, and (2) test these candidates for signatures of natural selection using population genetic statistics. We discovered nine putative inversion loci, which were supported by patterns of linkage disequilibrium, nucleotide diversity, and *F*_ST_ statistics. We investigated the relative age of genetic variation, and the distribution of genetic differentiation within and outside of loci of interest. These analyses, together with gene functional enrichment and mutational impact assessments, suggest that three of the nine loci show signatures of natural selection: one shows evidence of positive selection, while the other two show signatures of balancing selection. Overall, these results highlight that putative chromosomal inversions in *S. purpuratus* contribute to adaptive differentiation in the presence of high gene flow.

## RESULTS

### Nine Potential Inversions Discovered in *S. purpuratus*

We used the genomic dataset from Petak et al., (2023) to identify potential structural variants including cosmopolitan inversions across the range of the species. The original dataset contained 140 individuals, yet here we used 137 individuals after removal of 3 outlier individuals. Individuals were sequenced to an average depth of 6X coverage, and harbored ∼10 million SNPs across 21 chromosomes. The samples originated from seven locations, two in Oregon (Fogarty Creek [FOG] and Cape Blanco [CAP]), and five in California (Kibesilah Hill [KIB], Bodega Head [BOD], Terrace Point [TER], Lompoc Landing [LOM], and San Diego [SAN], from north to south; **Figure 1**). We applied the local Principal Component Analysis (PCA) method (Li and Ralph 2019) to each of the 21 chromosomes separately to find continuous genomic regions with PCA patterns different from the background. Results were compared across four different window sizes (500, 1000, 5,000, 10,000 SNPs; the results were robust across window sizes genome-wide (see **Figures S1, S2**).

**Figure 1.**
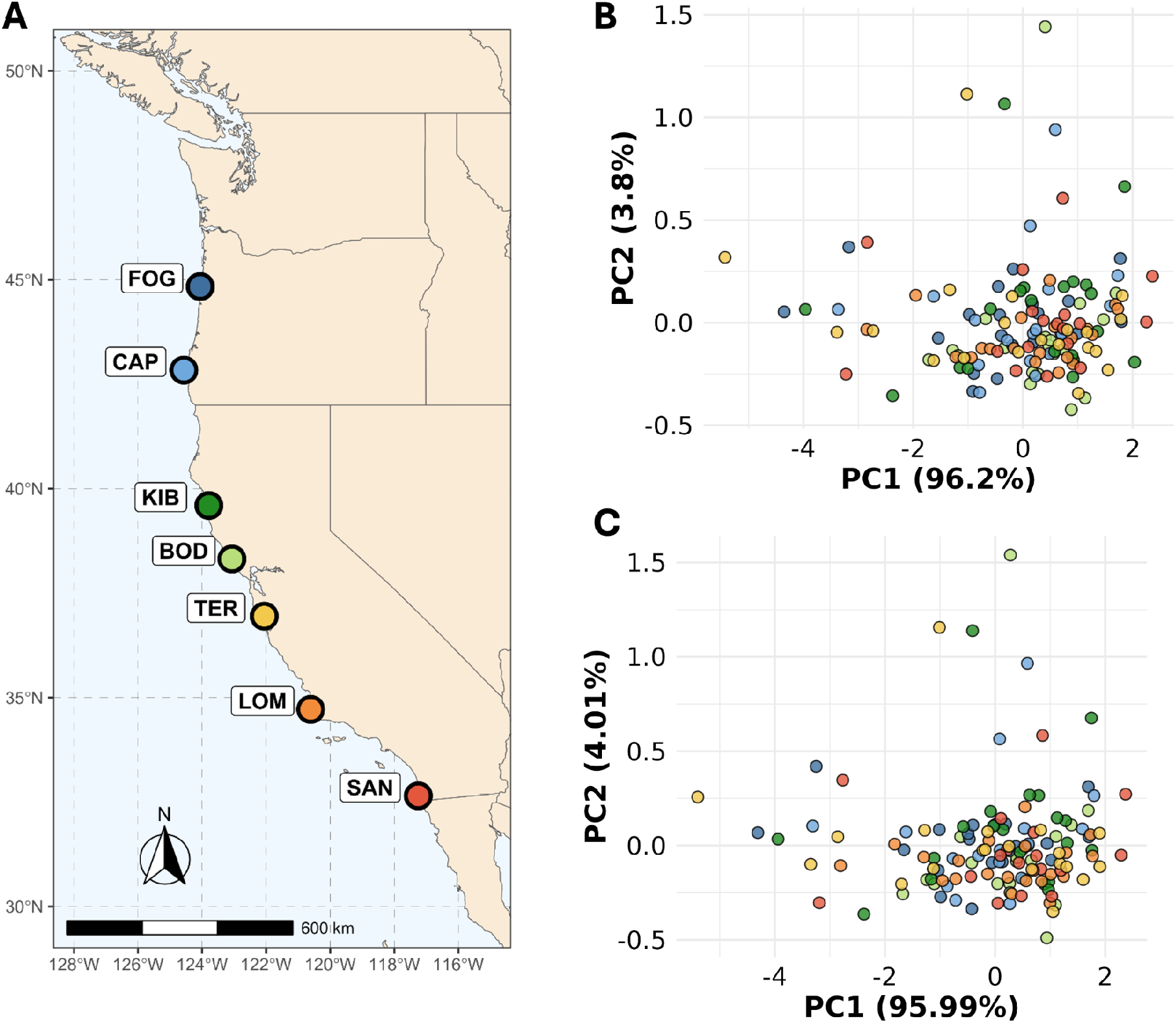
(a) Geographic distribution of samples (n=20 individual urchins per site) used in this study obtained from Petak et al., (2023). Principal Component Analyses (b) with (n= 8,139,929) and (c) without (n= 6,112,682) SNPs in putative inversions included. Both analyses excluded outlier loci to focus on neutral population structure.

We identified nine regions across eight chromosomes exhibiting consistently high absolute MDS values (in either of two dimensions) across all window sizes tested. These loci ranged in size from four million to 80 thousand base pairs long with a frequency of heterozygotes ranging from 15-60% (**Table 1**). The PCA clustered individuals into three distinct karyotypic groups (i.e., *locus 1, 3, 4, 8*, and *9*; **Figure 2**). For some of the loci, the three clusters were further divided along PC2 (i.e., *locus 2, 5, 6*, and *7*), indicating the presence of more haplotypes with three, or more, karyotypic alleles segregating in the population. See **Figure S3** for a more detailed exploration of these patterns. An additional parallel analysis similar to local PCA, sliding window PCA, showed areas of divergence between homozygote karyotypes in the same putative inversion regions (**Figure S4**). To confirm that the lack of population structure was not driven by the potential impact of structural variants obscuring geographic signatures (Galià-Camps et al., 2025), we ran PCAs with and without SNPs in putative inversion loci and found no difference in the pattern of population structure (**Figure 1B, C**) and conducted a pairwise analysis of the euclidean distances (**Figure S5**), and similarly found no difference between the distances within versus between populations for all SNPs (W = 4802274, p-value = 0.7519) and for all SNPs excluding those in putative inversions (W = 4801498, p-value = 0.745).

**Table 1:**
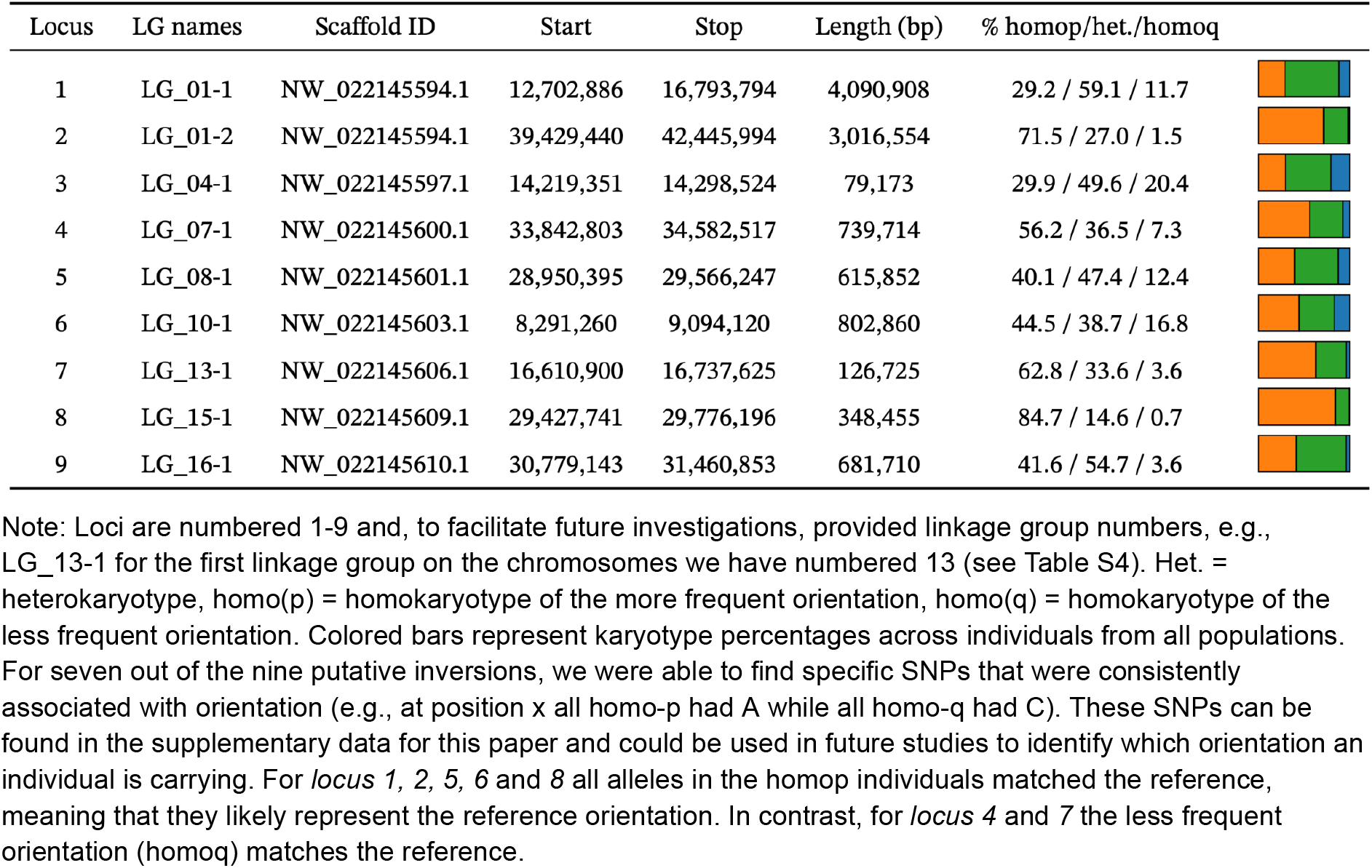
Genomic position, length, and percentage of individuals belonging to each karyotype group for the nine loci identified as putative inversion polymorphisms.

| Locus | LG names | Scaffold ID | Start | Stop | Length (bp) | % homop/het./homoq |
| --- | --- | --- | --- | --- | --- | --- |
| 1 | LG_01-1 | NW_022145594.1 | 12,702,886 | 16,793,794 | 4,090,908 | 29.2 / 59.1 / 11.7 |
| 2 | LG_01-2 | NW_022145594.1 | 39,429,440 | 42,445,994 | 3,016,554 | 71.5 / 27.0 / 1.5 |
| 3 | LG_04-1 | NW_022145597.1 | 14,219,351 | 14,298,524 | 79,173 | 29.9 / 49.6 / 20.4 |
| 4 | LG_07-1 | NW_022145600.1 | 33,842,803 | 34,582,517 | 739,714 | 56.2 / 36.5 / 7.3 |
| 5 | LG_08-1 | NW_022145601.1 | 28,950,395 | 29,566,247 | 615,852 | 40.1 / 47.4 / 12.4 |
| 6 | LG_10-1 | NW_022145603.1 | 8,291,260 | 9,094,120 | 802,860 | 44.5 / 38.7 / 16.8 |
| 7 | LG_13-1 | NW_022145606.1 | 16,610,900 | 16,737,625 | 126,725 | 62.8 / 33.6 / 3.6 |
| 8 | LG_15-1 | NW_022145609.1 | 29,427,741 | 29,776,196 | 348,455 | 84.7 / 14.6 / 0.7 |
| 9 | LG_16-1 | NW_022145610.1 | 30,779,143 | 31,460,853 | 681,710 | 41.6 / 54.7 / 3.6 |

Note: Loci are numbered 1-9 and, to facilitate future investigations, provided linkage group numbers, e.g., LG\_13-1 for the first linkage group on the chromosomes we have numbered 13 (see Table S4). Het. = heterokaryotype, homo(p) = homokaryotype of the more frequent orientation, homo(q) = homokaryotype of the less frequent orientation. Colored bars represent karyotype percentages across individuals from all populations. For seven out of the nine putative inversions, we were able to find specific SNPs that were consistently associated with orientation (e.g., at position x all homo-p had A while all homo-q had C). These SNPs can be found in the supplementary data for this paper and could be used in future studies to identify which orientation an individual is carrying. For *locus 1, 2, 5, 6* and *8* all alleles in the homop individuals matched the reference, meaning that they likely represent the reference orientation. In contrast, for *locus 4* and *7* the less frequent orientation (homoq) matches the reference.

**Figure 2.**
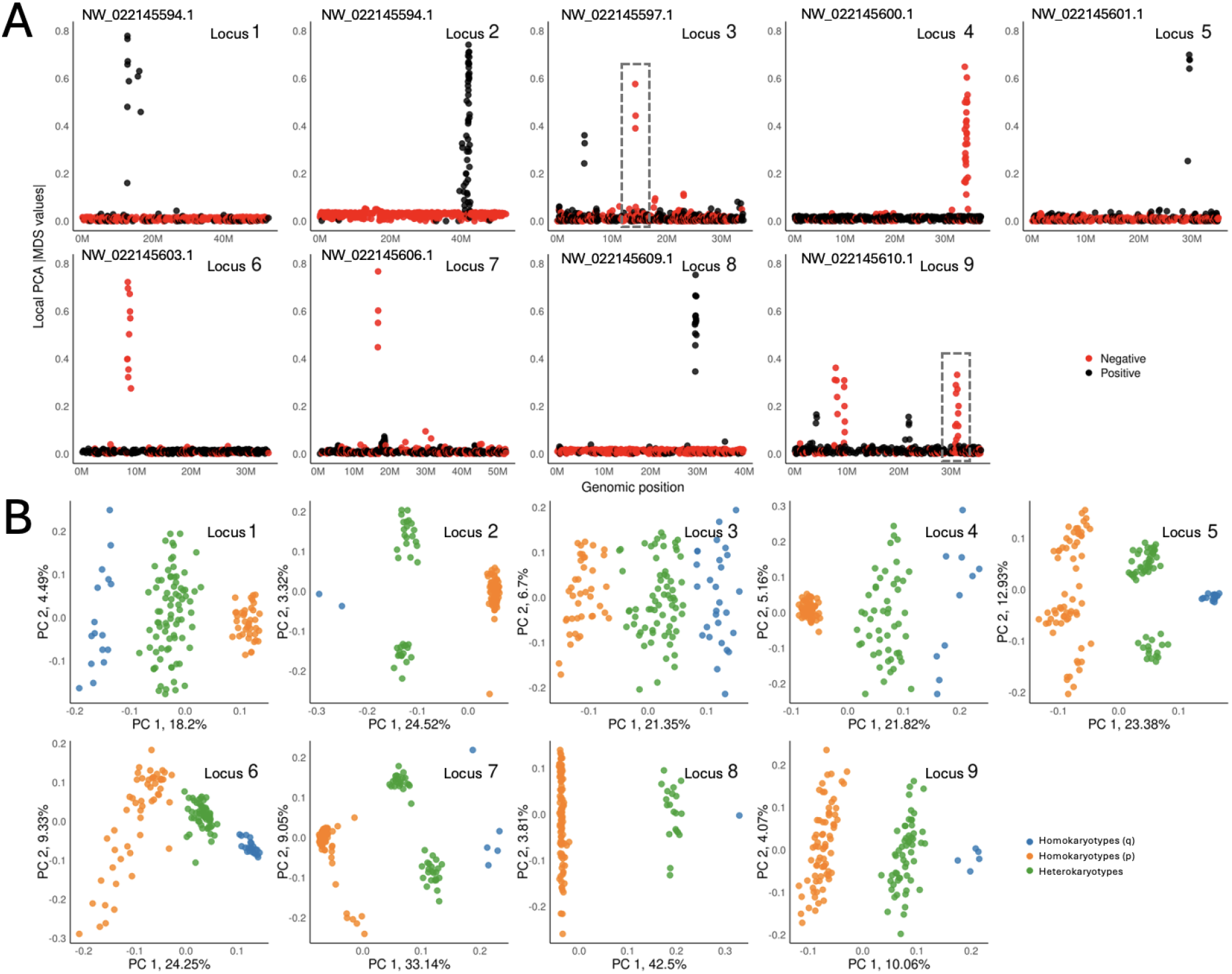
Nine regions with signatures of inversion polymorphisms. (A) The folded distribution (black: positive; red: negative) of MDS values along the whole chromosome (MDS1, except for 1, 3 and 9 MDS2). For chromosomes with multiple spikes, we added a dashed rectangle to indicate which of the MDS spikes correspond to the three-way clustering in panel B. (B) PCA plots for each locus computed using genotype data restricted to the region with the MDS spike. For these nine regions, PCA clustered individuals into three separate groups with three colors corresponding to the three karyotype groups: green = heterokaryotypes, blue = homokaryotypes (q), orange = homokaryotypes (p).

### The evolutionary properties of potential inversion polymorphisms in *S. purpuratus*

To assess whether the nine regions identified through local PCA exhibited classical evolutionary signatures of structural variants including inversion polymorphisms, we characterized a range of standard population genetic statistics across all nine loci. Since inversions suppress recombination, especially near breakpoints, we expect elevated linkage disequilibrium in these regions. In general, except for *locus 9*, we observed elevated levels of long-distance linkage disequilibrium (LD) relative to background LD, consistent with large blocks of suppressed recombination (**Figure 3**), a hallmark of inversions. Similarly, both homokaryotype groups, referred to here as “homokaryotype *p*” (i.e., major allele) and “homokaryotype *q*” (i.e., minor allele), showed elevated levels of genetic differentiation (measured as *F*_ST_; **Figure 4; Figure S6**). However, in four loci (i.e., *locus 2, 7, 8, and 9*), fewer than five homozygous inverted individuals were observed, which may reduce the statistical power to detect differentiation (**Table S1**). Among the nine loci examined, *locus 1, 3, 4 and 6* showed the strongest divergence at the putative inversion breakpoints. However, *locus 4* and *6* showed a clear hanging bridge pattern of *F*_ST_, whereas this pattern is less pronounced for *locus 1*, the longest of the loci (∼4 Mbp), as well as for *locus 2, 3* and *5*. Under the classic hanging bridge model (Berdan et al., 2023; Guerrero et al., 2012; Navarro et al., 2000, 1997), differentiation is expected to be strongest at breakpoints and to weaken toward the core, as secondary crossing over becomes more likely. In this context, the weaker *F*_ST_ and linkage pattern seen at *locus 1, 3* and *5* (**Figure 3**) may suggest complex histories of double crossovers, the presence of overlapping inversions, or alternatively, misassemblies in the reference genome. Indeed, the patterns at *locus 5* may comprise two overlapping inversions, a possibility partially supported by the large number of genotype clusters observed in PCA space in **Figure 2B**. We also examined nucleotide diversity inside and outside of each locus, but revealed no clear differences between the locus and the surrounding genomic region (**Figure S7**).

**Figure 3.**
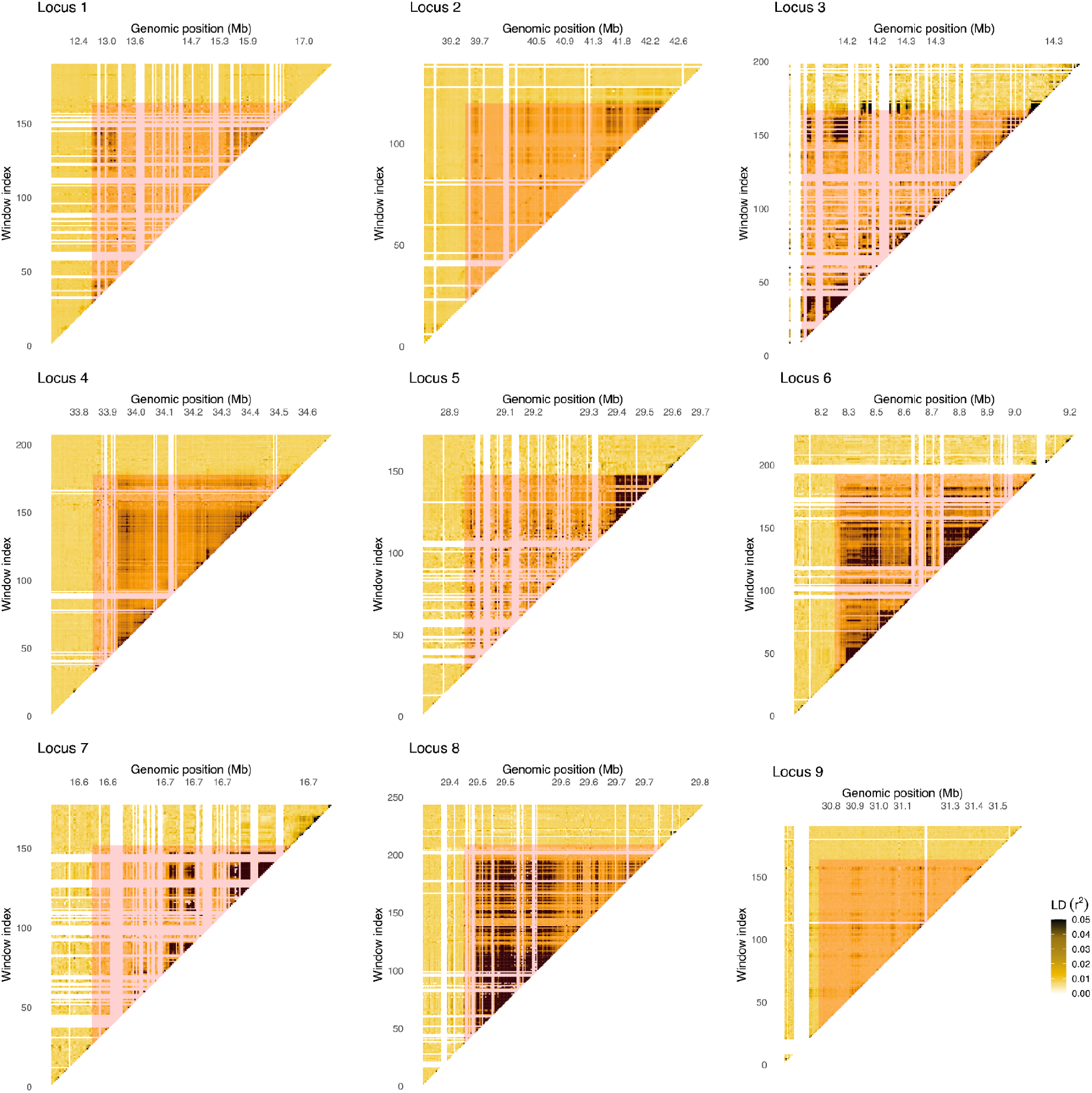
Linkage disequilibrium (LD) of *locus1-9*. LD value (*r*^2^) is coloured per SNP window from light yellow to black. Window sizes are 30,000 bp for *locus 1* and *2*, 500 bp for *locus 3*, 5000 bp for *locus 4-6* and *9*, 1000 bp for *locus 7*, and 2000 bp for *locus 8*. Putative inversion sites highlighted in red (Table 1). See Figure S6 for results of LD when only considering homokaryotypes.

**Figure 4.**
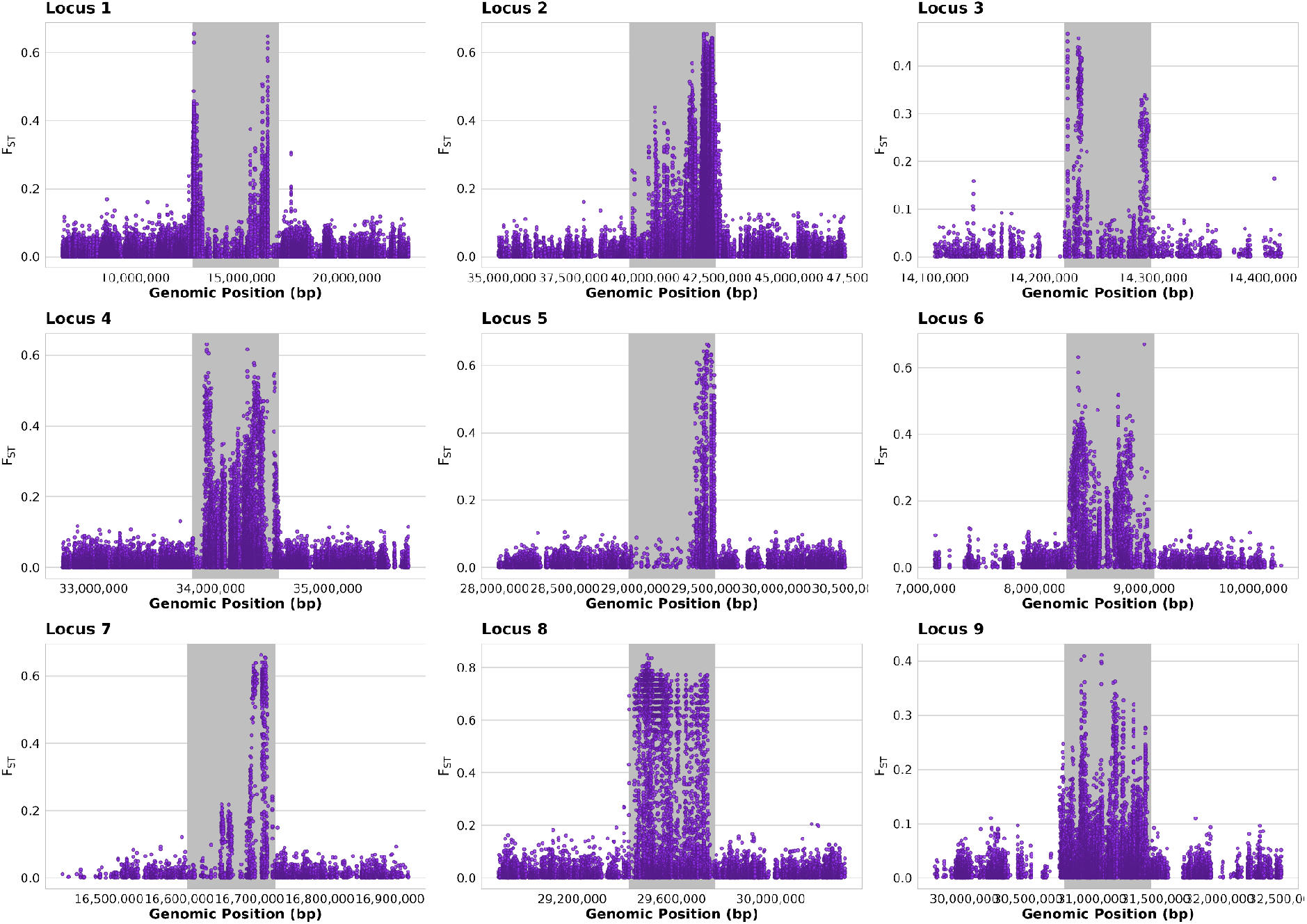
*F*_ST_ of heterozygotes for each locus showing 1.5x the inversion area above and below each locus region base. Grey area indicates the base of the putative inversion region.

Tajima’s D was also computed for each homokaryotype group individually across all loci (**Figure S8**). Notably, at locus 6 we observed elevated Tajima’s D in the homokaryotype-*p*, contrasted with a marked reduction in the alternative homokaryotype. This signature may indicate the action of an intra-karyotype selective sweep (see discussion). In addition, we estimated the age of mutations within and outside the locus regions. We also tested for a correlation between allele age and coverage and found no evidence of an association (*cor* = 0.00824, *P-value* = 0.0710). Generally, SNPs located within a locus were older than those outside except for *locus 6*, where SNPs inside the locus were comparatively younger (**Figure 5A**). Interestingly, *locus 1, 5*, and *7* appeared particularly old, with a time to the most recent common ancestor (T_MRCA_) ∼35% greater than the genome-wide average, exceeding that of all other chromosomes and locus regions. Furthermore, comparing the relative ages of mutations at the core versus at the breakpoints revealed no substantial differences in allele age, except for *locus 7* and *9*, where variants at the breakpoints appeared older than those at the core (*locus 7*: *P*-value = 0.033; *locus 9*: *P*-value = 0.070).

**Figure 5.**
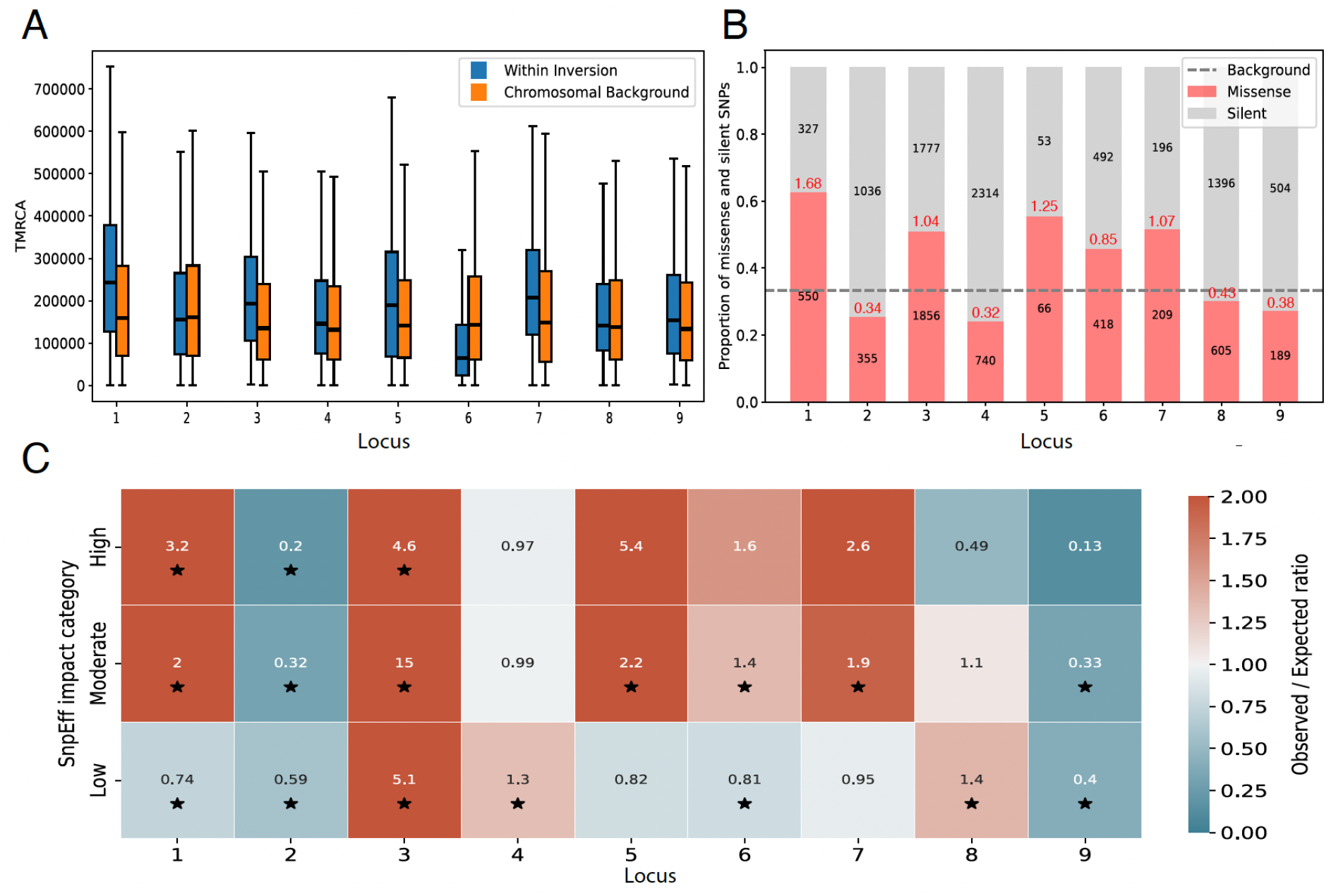
(A) Estimated Time to Most Recent Common Ancestor (T_MRCA_) for SNPs inside versus outside of each locus. All nine showed significant differences (Mann–Whitney U, *P*<0.0001). SNPs in *locus 2* and *locus 6* are younger, whereas for the rest of the seven loci SNPs are older than expected based on the background. (B) Number of SNPs with missense or silent effect in each locus. Red numbers show the missense / silent ratio. The average missense / silent ratio across the chromosomes outside of loci was 0.5, meaning that the expectation is that there were twice as many silent than amino acid changing SNPs (gray dashed line). *Locus 1* had a significant increase in missense SNPs, in contrast, the majority of SNPs in *locus 2* were silent. All ratios are significantly different from expectation based on two-sided binomial tests with Bonferroni correction for multiple testing. (C) Ratio of observed SNPs with high, moderate, or low impact to the expected number based on the chromosomal background (color and value). SNP impacts were categorized using SnpEff. Red: more observed than expected, blue: less observed than expected. Stars indicate statistical significance based on two-sided binomial tests with Bonferroni correction for multiple testing.

The frequency of the individuals belonging to different karyotype groups differed from the expectation based on the Hardy-Weinberg Equilibrium only in *locus 1* (*χ*^2^ = 6.6, *P-value* = 0.036). There were more heterokaryotypes (81 observed vs 66.39 expected) and fewer homokaryotypes in either orientation (16 observed vs 23.3 expected and 40 observed vs 47.3 expected) than expected. In addition, heterokaryotype frequency increased sharply south of Kibesillah Hill (see **Figure S9** for correlation with latitude for all nine putative inversions), with a significant difference between the three northern and four southern populations (*P-value* = 0.0015). Heterokaryotypes at Locus 5 were also correlated with latitude, however with heterokaryotypes more common in the north (R^2^ = 0.64, *P-value* = 0.0316; **Figure S3; S9**).

We examined genome annotations to assess the missense-to-silent mutation ratio within the identified loci, relative to the genomic background. We found that this ratio varied substantially across the loci of interest (mean = 0.80 ± 0.48; **Figure 5B**), in contrast to the relatively consistent background ratio observed across chromosomes (mean = 0.50 ± 0.08). Using the SnpEff package, we categorized SNPs as high impact – likely to have a large effect on protein function such as truncation or loss of function (e.g., stop gain and frameshift variants), moderate impact – less disruptive SNPs that change protein effectiveness (e.g., missense or inframe deletion), and low impact – unlikely to change protein function (e.g., synonymous variants). Like above, the proportion of SNPs with high, moderate, or low impact within loci varied across putative inversions with *locus 1, 3, 5*, and *7* showing overrepresentation and *locus 2* showing underrepresentation of moderate and high impact SNPs (**Figure 5C**).

### Functional Gene Ontology Analyses Across Loci

Across the nine loci examined, two had enrichment of specific functional classes of genes. *Locus 1*, the largest putative inversion (∼4 Mbp), contained 72 genes, with only the cell-cell adhesion category enriched after correction for multiple-testing. In contrast, *locus 2*, the second largest putative inversion (∼3 Mbp), contained a similar number of genes, 76, but had many enriched functional classes of genes, notably regulation of cellular response to hypoxia (GO:1900037) and regulation of telomere maintenance via telomere lengthening (GO:1904356). Two genes within the hypoxia category, LOC592339 and LOC105441271, both members of the *Hsp70* chaperone family, carried multiple variants including nonsynonymous and UTR changes.

None of the other loci had functional enrichment. However, *locus 6* showed a small *F*_ST_ spike adjacent to its right breakpoint (**Figure 4**), coinciding with clusters of 5S ribosomal RNA genes, suggesting a repetitive DNA region; excluding these, 21 genes mapped to the area, and are all associated with the carnitine biosynthetic process GO term (GO:0045329). *Locus 7* spanned seven genes, including LOC763759, annotated as zinc transporter ZIP14 (*Sp-Slc39a8*) which is involved in bicarbonate transport during biomineralization (Jenkitkasemwong et al., 2012).

### ootprints of Natural Selection at Potential Inversion Polymorphisms

To evaluate whether any of the nine loci of interest were subject to natural selection, we performed a genome-wide scan using the Bayesian software *BayPass* to estimate the *X*^*T*^*X* statistic while accounting for population structure (Günther and Coop, 2013). An elevated *X*^*T*^*X* value relative to the genomic background suggests positive selection, whereas a reduced *X*^*T*^*X* suggests a signature of balancing selection. Although *X*^*T*^*X* is calculated on a per-locus basis, we complemented this analysis with a local score approach (i.e., a “Lindley Score”; Bonhomme et al., 2010), which scans the genome for windows enriched in outlier *X*^*T*^*X* values based *ξ* = 3 where only SNPs with *X*^*T*^*X P-values* < 0.0001 contributed to the score. Regions showing significant enrichment of outliers were considered candidate loci evolving under natural selection. Our *BayPass* analysis revealed 351 significant peaks which included 2769 outlier SNPs. Although a comprehensive analysis of these peaks lies beyond the scope of this study, we examined the extent to which these signals of selection overlapped with our candidate chromosomal inversions. Indeed, mapping putative inversions to outlier regions likely under selection (i.e., peaks), we found overlap at *locus 2* (8 SNPs) *locus 6* (128 SNPs), *locus 7* (14 SNPs), and *locus 8* (105 SNPs; **Figures 6A, S10**). Within these loci, *locus 6, locus 7* and *locus 8* were significantly enriched for *X*^*T*^*X* outliers (**Figure 6B; Table S2**). Estimating the relative age of *X*^*T*^*X* outliers in comparison to the inferred boundaries (i.e., the putative breakpoints of each locus) showed that, for *locus 7*, the outliers were similar in age to both the breakpoints and the rest of the putative inversion. In contrast, for *locus 8*, the *X*^*T*^*X* outliers appeared older than both the breakpoints and other internal regions of the locus (*P*-value = 0.00165; **Figure 6D**) and for *locus 6*, the *X*^*T*^*X* outliers appeared younger than the breakpoints (*P*-value = 0.0124; **Figure 6D**).

**Figure 6.**
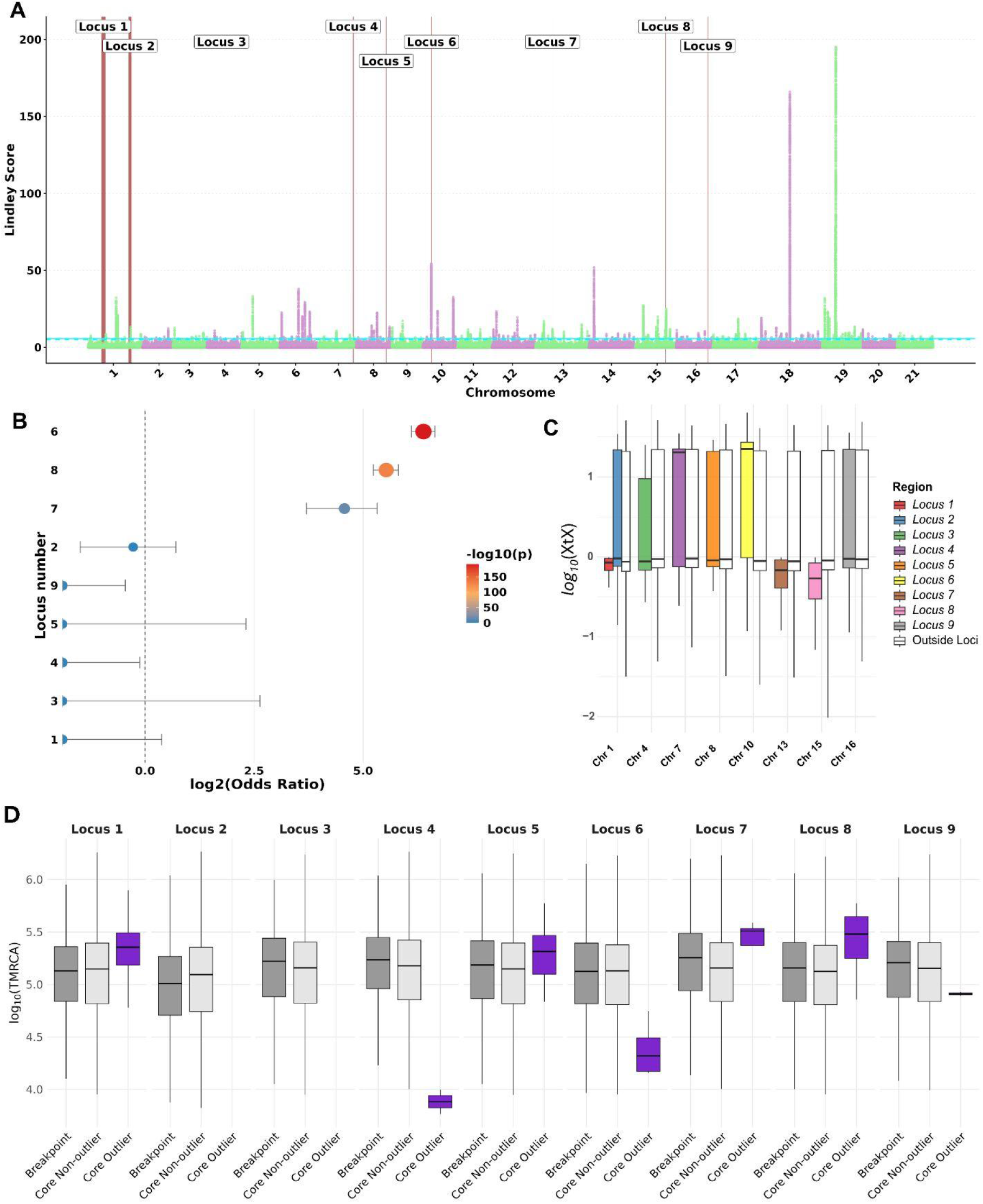
Markers of selection across loci containing putative inversions. (a) Manhattan plot of SNPs displaying Lindley score values calculated from *localscore* grouped by chromosome with putative inversion loci highlighted in red and numbered. Horizontal blue lines indicate Lindley significance threshold calculated in localscore, dotted indicate lower threshold and smooth indicates upper threshold based on chromosome by chromosome calculations of Lindley. (b) Fisher’s Exact Test odds ratios for enrichment for *X*^*T*^*X* outliers across loci. Size of dot indicates -log_10_ *P-*value and error bar indicates ± SD. (c) Distribution of log_10_(*X*^*T*^*X*) within (colored) versus outside (white) each locus of interest per chromosome. (d) Distribution of log_10_ Time to the Most Recent Common Ancestor (T_MRCA_) for breakpoint (dark grey), core non-outliers for *X*^*T*^*X* (light grey), and outliers for *X*^*T*^*X* (purple) per locus of interest.

Additionally, we found the *X*^*T*^*X* scores were lower in *locus 7* and *locus 8* than SNPs outside the loci within that chromosome (*P-value* < 0.001; **Figure 5C, Table S3**) consistent with signals of balancing selection, while *X*^*T*^*X* scores in *locus 6* were higher than scores outside the locus within that chromosome suggesting a strong signal of directional selection (*P-value* < 0.001; **Figure 5C, Table S3**). While the annotations of all genes are currently uncharacterized, BLASTp analysis of protein sequences revealed that 43 (8% of SNPs in *locus 8*) were located in a gene homologous to *apolipoprotein B* (Query Coverage = 59%, Percent Identity = 94.13%, *e*-value = 1.0 × 10^−300^, species = *Mesocentrotus nudus*), a gene implicated in lipid transport (Yuhi et al., 2022).

## DISCUSSION

Here, we present a genome-wide assessment of loci that are potentially inversion polymorphisms in the model sea urchin, *Strongylocentrotus purpuratus*. Whole genome sequences from 137 diploid individuals sampled from seven sites along the west coast of North America revealed patterns of suppressed recombination suggestive of inversion polymorphisms. Among the nine loci identified as putative inversions, most showed long-range LD among their putative breakpoints and the characteristic *F*_ST_ “bridge pattern,” where breakpoints show elevated differentiation relative to the core (Guerrero et al., 2012) which are striking patterns given the rapid genome-wide LD decay in purple urchins (Brennan et al., 2019; Petak et al., 2023), lending further support to the presence of cosmopolitan structural variants.

Patterns of genetic variation captured in the PCA (**Figure 2B**) provide insight into the haplotypic structures that underlie these putative inversions. For example, *locus 1, 3, 4, 8*, and *9* each exhibit three genotype clusters along PC1, consistent with two homozygous classes (standard and inverted) and a heterozygous class. *Locus 6* also shows three well-defined clusters; however, it also appears to harbor footprints of an intra-karyotypic selective sweep (see below). In contrast, *locus 2, 5*, and *7* display multiple clusters distributed across both PCs 1 and 2, a pattern that may be consistent with either the evolution of multiple, highly diverged, inverted haplotypes (Berdan et al., 2021), or the presence of overlapping inversions. Notably, the triangular clustering pattern in *locus 5* resembles the overlapping inversion structure described for the *LGC6*.*1/LGC6*.*2* inversions in *Littorina saxatilis* (Faria et al., 2019).

### Convergent evidence identifies three strong candidates as putatively adaptive inversions

Of the nine putative inversion polymorphisms, three, *locus 6, 7*, and *8*, showed evidence of natural selection, as suggested by enrichment of *X*^*T*^*X* outliers. Interestingly, these three loci exhibit markedly different age estimates relative to the genome-wide background. *Locus 7* appears older than the background, locus 8 is comparable in age to the background, while *locus 6* is substantially younger. These estimates provide insight into the potential processes giving rise to these adaptive inversions. Two main models have been proposed to explain the origin of adaptive inversions: “gain” and “capture” (Schaal et al., 2022). Under the “capture” model, inversions capture pre-existing (i.e., older than the inversion) combinations of locally adapted alleles, and suppressed recombination among these beneficial variants reduces the chance of recombination load. This has been described in systems such as Atlantic cod (Kirubakaran et al., 2016) and *Mimulus* (Coughlan and Willis, 2019). In contrast, the “gain” model posits that inversions arise as neutral variants and subsequently accumulate adaptive mutations (i.e., adaptive alleles should be younger than the inversion itself). An example is the inversion underlying alternative mating morphs in *Philomachus pugnax* (Lamichhaney et al., 2016). In our data, *locus 6* and *7* show a pattern in which mutations in the core region, including *X*^*T*^*X* outliers, are significantly younger than those near the breakpoints. This pattern is consistent with the gain model. In contrast, at *locus 8* we observe that *X*^*T*^*X* outliers are significantly older than the breakpoints, a pattern consistent with the capture hypothesis. An important caveat is that our estimation of allele ages were derived from a statistically phased SNP panel using coalescent modeling. A known drawback of this approach is that low coverage and phasing errors can artificially reduce the estimated age mutations (Albers and McVean, 2020). Thus, age estimates should be interpreted with caution, although we expect this issue to have a systematic effect across the dataset. Consistent with this, we observed no correlation between allele age and coverage.

Functional annotations suggest potential biological relevance of these inversions. For example, *locus 6* is enriched for a GO term associated with carnitine biosynthetic processes, which may be important for energetic efficiency (Bremer, 1990). *Locus 7* was significantly enriched for zinc transporter (ZIP14) functions, which is hypothesised to be involved in biomineralization (Hu and Jiang, 2024; Jenkitkasemwong et al., 2012). This may be relevant to adaptation across pH gradients (e.g., Petak et al., 2023). *Locus 8*, was enriched for three GO terms associated with lipid transportation and localization, functions which may be key for growth and development (Li et al., 2021). In addition, in our adaptive differentiation scan (i.e., *X*^*T*^*X*) we identified a homolog of *apolipoprotein B*, inside *locus 8*, as a candidate gene under selection. Assuming the homologous function is retained, this gene could be ecologically important as it is associated with the transport of key nutrients to the eggs from the digestive system via the coelomic fluid (Yuhi et al., 2022). Despite these interesting signals, extended linkage disequilibrium from suppressed recombination in inversions complicates causal variant identification, especially near breakpoints. As such, future work should focus on karyotype validation and fitness assays under relevant environmental stressors (e.g., pH, temperature, or oxygen variation).

### Putative adaptive inversions show distinct evolutionary histories

Although all three candidate adaptive inversions, *locus 6, 7*, and *8*, were enriched for *X*^*T*^*X* outliers, they displayed different patterns of genetic variation across our analyses. For example, we observed elevated levels of missense mutations at *locus 6* and *7*, a pattern that likely represents the accumulation of genetic load. Both empirical and simulation studies (Berdan et al., 2021; Betancourt et al., 2009; Campos et al., 2014; Charlesworth, 1996) suggest that this buildup of deleterious, likely recessive, variation, is a common and potentially inevitable consequence of reduced recombination that ultimately leads to reduced fitness. One proposed solution to the problem of genetic load in inversions is the evolution of haplotype substructuring, whereby multiple divergent genetic backgrounds arise from the original inverted arrangement to mitigate fitness loss across haplotypes (Berdan et al., 2021; Charlesworth and Charlesworth, 1997). Indeed, our PCA suggests this may be occurring at *locus 7*, where complex genotypic clustering in PC space is consistent with diverged karyotypes (**Figure 2B**). Furthermore, the weak population structure in urchins (driven by high gene flow), suggests that populations from California to Oregon behave largely as a single metapopulation. This pattern is compatible with theoretical expectations for how haplotype substructuring may emerge (Berdan et al., 2021). In the case of *locus 6*, the putative inversion appears to be undergoing an intra-karyotype sweep (Charlesworth and Charlesworth, 1997; Nunez et al., 2024), as evidenced by the extended region of elevated Tajima’s D within the homokarytype-*q* group inside the inversion (**Figure S8**). This locus also shows the strongest enrichment of *X*^*T*^*X* outliers and is the youngest among the loci. In this scenario, genetic load may be mitigated through the wholesale replacement of genetic backgrounds.

Interestingly, *locus 8* showed no enrichment of missense mutations. This is unexpected, given the theoretical expectations of reduced purifying selection in inversions (Berdan et al., 2021; Crumpacker and Salceda, 1969; Dobzhansky et al., 1963). One possibility, consistent with our interpretation of the T_MRCA_ data, is that *locus* 8 captured ancient high fitness haplotypes experiencing strong selection. Another possibility is that the inverted karyotype has already accumulated an intolerable burden of recessive load, rendering it fully inviable in the homozygous state. However, it may remain viable in heterozygotes due to associative overdominance (Berdan et al., 2021). This scenario could explain why the inversion is predominantly observed either as a homozygote of one class or as a heterozygote, while alternative homozygous are extremely rare (**Figure S8**).

One important caveat to consider, is that these hypotheses assume that mutations inside the inversion are the drivers of fitness differentials. An alternative possibility is that these inversions could exert an effect through breakpoint-mediated regulation of a nearby gene, disrupting *cis*- or *trans*-regulatory architecture, with missense variation playing little or no role (Harewood and Fraser, 2014; Lavington and Kern, 2017). Ultimately, functional follow-up experiments will be required to disentangle the mechanisms underlying these loci.

### Limitations and alternative explanations for inferred inversions

A key limitation of our study is that SNP-based population genomic analyses from paired-end short reads are indirect, relying on detecting localized reductions in recombination from allele frequency patterns. While most loci identified show patterns consistent with inversions, several alternative explanations cannot be excluded, such as genome assembly errors and other causes of low recombination, such as traces of introgression from other species or as of yet unidentified sex-determining regions. For example, theoretical and empirical work has shown that inversions are unlikely to persist in populations as neutral polymorphisms (reviewed in Berdan et al., 2023). Why, then, do we only see signatures of selection in only three of the nine putative inversions? Some may not be real inversions, but rather assembly artifacts or otherwise regions of reduced recombination (e.g., incomplete sweeps). Indeed, local three-way clustering suggests the presence of two diverged haplotypes within populations (i.e., a common consequence of inversion polymorphism), however, similar patterns can also arise from other processes, such as natural selection acting on linked alleles (Li & Ralph, 2019). Another possibility is that these loci experience selection at spatial or temporal scales not captured in the sampling scheme of Petak et al. (2023), which we rely on here. For example, our dataset spans samples from California and Oregon, whereas the species’ range extends from Alaska to Baja California, Mexico (Ebert, 2010); consequently, important selective regimes across the broader geographic range may be missed, including spatially heterogeneous pressures such as barren versus kelp forest habitats (Dayton et al., 1992; Filbee-Dexter and Scheibling, 2014; Pearse, 2006). Depth represents an additional uncharacterized axis of ecological variation which has been shown to be important in other work on *S. purpuratus* (Rumberger et al., 2025). Given that all samples were derived from coastal adults, selective pressures associated with vertically structured oceanographic conditions are likely undersampled. Finally, interannual environmental variation may drive temporally fluctuating selection. Although urchins are long-lived, their habitats experience multi-decadal cycles, such as *El Niño* and *La Niña* events, which could impose episodic selection not captured in our dataset. Further sampling, long-read sequencing, and experimental work will be necessary to confirm whether these loci are indeed inversion polymorphisms, as well as to determine their functional roles and histories of selection.

## Conclusion

We identified nine genomic regions in the purple sea urchin that exhibit reduced recombination and strong linkage, likely corresponding to highly polymorphic inversions. For some of our candidate loci (e.g., *locus 6*) we found robust evidence supporting their classification as putative adaptive inversions. In contrast, other loci (e.g., *locus 5*, despite the latitudinal cline) remain inconclusive and require further study. Indeed, future studies comparing gene expression and phenotypic differences between inversion orientations, as well as experiments testing fitness effects under varied environmental stressors, will be essential to uncover the functional significance of these loci. Nevertheless, these findings broaden our understanding of genome evolution in natural populations and add to the genomic repertoire of *S. purpuratus*, a key model system in developmental and evolutionary biology.

## MATERIALS AND METHODS

### Data acquisition and genome subsampling

Sequencing reads for all 140 *S. purpuratus* individuals were downloaded from NCBI Sequence Read Archive BioProject accession number PRJNA674711. See (Petak et al., 2023) for detail on sample collection and sequencing. The reads were mapped to Spur v. 5.0 (scaffold N50 ∼37 Mbp), the same way as in (Petak et al., 2023). The reference genome Spur v5.0 consists of 870 scaffolds. However, sorting all of the scaffolds by size, it is clear that 21 scaffolds are much larger than the rest. The largest 21 scaffolds all have length greater than ∼30Mb and together they make up 89.87% of the data in the assembly. We focused our analysis on these scaffolds and since *S. purpuratus* is known to have 21 chromosomes (Eno et al., 2009), here, we refer to these scaffolds as chromosomes (see **Table S4** to link scaffold ID with chromosome numbers). Next, the bcftools (v. 1.10.2) mpileup algorithm was used to call variants for the largest 21 scaffolds (more detail on this in the Results section) (Danecek et al., 2021). The output bcf files were filtered by the following criteria: posterior genotype probability (QUAL >= 40), read depth across all individuals (DP > 560), Mann-Whitney U test of Mapping Quality Bias (MQB > -3), Mann-Whitney U test of Read Position Bias (-3 < RPB < 3) and total number of alleles in called genotypes (AN > 238). Variants with allele counts of less than 14 were discarded (5% MAF of 137 diploid genomes) and only biallelic single nucleotide polymorphisms (SNPs) were kept. Lastly, all SNPs that fell into repetitive regions according to the dataset published by (Panyushev et al., 2021) using RepeatModeler 2.0.1 were excluded from the dataset. We found the same three outlier individuals as (Petak et al., 2023), as many of the windows clustered them separately from the rest of the individuals along PC1. Thus, we excluded these three individuals from further analysis.

### Finding putative inversions

We used the lostruct package in R to perform local PCA along the genome (Li and Ralph, 2019). The local PCA method of identifying genomic regions with population structure different from the background consists of the following steps. First, the genome of interest is divided into fixed sized windows (here we used a certain number of SNPs, but the results were unchanged when defining windows according to the number of base pairs). All chromosome-specific local PCA analyses were repeated for four different window sizes, 500, 1000, 5000 and 10,000 SNPs. Then, for each window a principal component analysis is performed using the SNPs located within the window. Each window was then compared to every other window based on the first N principal components (two by convention) using a multidimensional scaling (MDS). Windows that cluster together in the MDS are windows where the PCAs for those windows are similar. For example, if there is a large genomic region where there is population structure, e.g., SNPs cluster the individuals by sampling location on the PCA, windows spanning this region will be closer together on the MDS plot than windows from a genomic region where individuals from different sampling locations overlap on the PCA. In our case, we are looking for population structure indicative of inversions. Inversions suppress recombination and therefore the two orientations independently accumulate genetic variation over time resulting in three-way clustering on the PCA based on the orientation of the inversion the individuals are carrying (two groups of homozygotes and one group of heterozygotes).

In this study, all MDS plots displayed the distribution of windows either in an L or a T shape (**Figure S2**), spreading windows along only one of two dimensions. This meant that we could plot the two MDS dimension values separately along the genome to find regions with a continuous series of outlier windows (i.e., windows where the individuals were clustered on the PCA the most different compared to the background). Three-way clustering was determined visually for each MDS outlier region on each chromosome, followed by analytical confirmation by the elbow method following Scikit-learn’s (Pedregosa et al., 2011) (v.1.4.2) agglomerative clustering with different k-values. A specific MDS outlier region was selected for further analysis if 1) the windows with MDS spikes were consecutive and 2) the elbow method showed clear three-way clustering. Both of these criteria needed to be met consistently across the four window sizes for the region to be selected.

The putative breakpoints were determined based on the location of the MDS spikes. The left breakpoint was determined to be the start position of the left-most window of the MDS spike, while the right breakpoint was determined to be the end position of the right-most window of the MDS spike. This was done using the smallest window size (500 SNPs). This process was automated using Scikit-learn’s K-means clustering of the window positions above the |0.1| MDS value threshold. These putative breakpoints were then supported by the *F*_ST_ and linkage disequilibrium results. We also generated nucleotide diversity values across the genome using Pixy v. 2.0.0 (Korunes and Samuk, 2021) using 1000 bp windows and default settings. Additionally, we used winPCA v. 1.2.1 (Blumer et al., 2025) to perform a sliding windowed PCA across loci regions, and we plotted PC1 per homokaryotype per loci.

### Population Genetic Statistics and Functional Enrichment

Each loci of interest (i.e., putative inversion) was extracted from the original filtered vcf file. These files were then converted to the gds file format to be used as an input to a principal component analysis (snpgdsPCA R package SNPRelate, (Zheng et al., 2012)). Based on the first two principal components, individuals were assigned to three groups using agglomerative clustering (Scikit-learn). Individuals were genotyped as heterokaryotypes if they belonged to the middle group along PC1, and as homokaryotypes if they belonged to groups on the right or left of the middle group. Following the standard convention of p for the more frequent and q for the less frequent allele, the homokaryotype group with more individuals was named homo-p and the group with less individuals as homo-q.

For each putative inversion deviation from the Hardy-Weinberg equilibrium was determined using a chi-squared test, and the correlation between genotype and allele frequency with geographical latitude (from (Petak et al., 2023)) was assessed using simple linear regression. Per site *F*_ST_ between the homokaryotype groups was calculated using vcftools (v.0.1.14) (Danecek et al., 2011) “weir-fst-pop” flag which estimates *F*_ST_ in accordance to (Weir and Cockerham, 1984). Vcftools was also used to calculate π diversity using the “window-pi” flag, and Tajima’s D using the “TajimaD” flag, both with a window size of 100bp. Linkage disequilibrium was calculated from the gds files in windows (window size dependent on the size of the region of interest such that around 200 windows are compared) using the SNPRelate snpgdsLDMat function with “corr” option for returning the correlation coefficient. To check whether genes located within the putative inversions were significantly enriched for specific GO categories compared to all annotated genes across the genome, we used a standard GO enrichment analysis. GO enrichment was conducted using GO terms retrieved from UniProt (The UniProt Consortium, 2025), followed by enrichment calculation with topGO (Alexa and Rahnenführer, 2022) and the GO enrichment tool on https://geneontology.org/ using the Bonferroni correction for multiple testing (Aleksander et al., 2023; Ashburner et al., 2000; Thomas et al., 2022). Finally, the SnpEff software (Cingolani et al., 2012) was used to categorize the effects of SNPs inside and outside of loci of interest (e.g., nonsynonymous/synonymous coding variants). A two-sided binomial test was used to determine the significance of differences between the number SNPs inside the loci with certain effects compared to the expectation given SNPs outside of the loci. Expectation was scaled by the frequency of variants in intergenic regions to account for potential biases in gene density.

### Allele Age Estimates

To find the common ancestor of each inversion, T_MRCA_ was computed using the Genealogical Estimation of Variant Age (GEVA; Albers and McVean, 2020). First, the variant call files were phased using the statistical phasing software shapit5 (Hofmeister et al., 2023). Since *S. purpuratus* has no known mutation rate, recombination rate, or effective population size, we used parameters from *Drosophila melanogaster*. Thus, the T_MRCA_ values calculated here should not be converted into years, rather, they are meant to reflect relative differences in ages between SNPs. Mann–Whitney U tests were used to compare T_MRCA_ values inside and outside of loci.

### Identifying regions under selection

To identify regions experiencing selection we ran an analysis with *BayPass* v. 2.41. Due to the lack of population structure (Petak et al., 2023) we ran the core *BayPass* model, randomly sampling and mixing the genome over 1000 chunks, then reassembling them in the correct order. *BayPass* was set to seven populations, with the rest of the settings as default. Peaks were then identified from the *X*^*T*^*X* output using the package *localscore* (Robelin et al., 2025). Additionally, we ran a Fisher’s exact test to identify loci that were significantly enriched for *X*^*T*^*X* outliers found from the Baypass analysis.

We chose BayPass because marine species such as sea urchins typically exhibit high gene flow and weak population structure (Petak et al., 2023), which complicates the detection of selection due to miscalibrations against demographic expectations (Lotterhos and Whitlock, 2014). *BayPass* addresses this issue by incorporating a genetic relatedness matrix (i.e., and Ω matrix) to account for population structure (Bonhomme et al., 2010; Gautier, 2015). Consistent with previous work (see Petak et al., 2023) we treated our seven populations along the west coast as samples from a well mixed metapopulation. In this context, we interpreted locus specific *X*^*T*^*X* values as relative to the model assumptions such as low *X*^*T*^*X* outliers are consistent with balancing selection, whereas high *X*^*T*^*X* outliers are consistent with positive selection across the heterogeneous landscape.

## Data Availability

All analyses in this study were based on publicly available datasets. Annotated VCF at https://doi.org/10.5281/zenodo.18613723. All code is available in GitHub at: https://github.com/Cpetak/Urchin_inversions

## Acknowledgements

This work was supported by the National Science Foundation award IOS 1943396 to M.H.P. This work was also supported by start-up funds from the University of Vermont to J.C.B.N. D.E.S salary was supported by the Office of the Vice President for Research at the University of Vermont. The authors acknowledge the Vermont Advanced Computing Center (VACC; https://www.uvm.edu/vacc) for providing computational resources that contributed to this publication.

