## Supplemental Information for "Evidence of Adaptation in Structural Variants among Wild Populations of the purple sea urchin, *Strongylocentrotus purpuratus*"

***Page Contents***

**S2 Supplementary text**

**S3 Table S1; Table S2**

**S4 Table S3; Table S4**

**S5 Figure S1**

**S6 Figure S2**

**S7 Figure S3**

**S8 Figure S4**

**S9 Figure S5**

**S10 Figure S6**

**S11 Figure S7**

**S12 Figure S8**

**S13 Figure S9**

**S14 Figure S10**

### Supplementary text

#### *Locus 5 – Six-way clustering*

Interestingly, while PC1 separated the individuals three-way PC2 further divided the groups in this inversion (Figure 1B). There were a total of 6 groups on the PCA, 3 on the left, 2 in the middle and one on the right. If there are 3 alternative combinations of SNPs (i.e., haplotypes) in the population, say A, B and C, then there are 6 possible combinations of haplotypes in diploid individuals - AA, AB, AC, BB, BC and CC. Since there are 3 groups along PC1, this might mean that individuals with AA, AB, and AC are grouped together (orange), BB and BC form the second group (green), and CC is its own group on the right (blue) on Figure 1B. Furthermore, the fact that 3 of the groups have half the number of individuals than the other 3 groups (11, 16, 17 vs 26, 28, 39) support this hypothesis since with 3 haplotypes there are three possible homozygotes and three heterozygotes, the latter being twice as likely to appear. Indeed, one of the smaller groups is the group on the right hypothesized to be CC (17 individuals), and we can hypothesize following the same reasoning that the bottom green group is BB (16) and the bottom orange group is AA (11). Consequently, the top green group is BC (39), and since PC2 separates by the second haplotype - and BC is above BB - the top orange group is AC (28) and the middle orange group is AB (26).

The number of homokaryotypes and heterokaryotypes was not significantly different from the HWE neutral expectation (neither assuming two (p=0.61) or three (p=0.98) haplotypes).
There was a significant correlation between the frequency of heterokaryotypes (assuming two haplotypes) and latitude, $r^2 = 0.64$, $p = 0.0316$, with more heterokaryotypes in the North (see Figure S5 below). No additional pattern emerged from looking at the correlation between the 6 genotypes and latitude, or the three haplotypes and latitude. There seems to be an increase of CC in the South, though the linear regression was not significant (p=0.14).

#### *Locus 7 – Six-way clustering*

Like in *locus 5*, interestingly, PC2 (9.05%) further divided the three groups into two, resulting in a total of 6 groups in this inversion (Figure 1B). The relative position of these 6 groups is very different from the one we saw in *locus 5*. The distribution of the number of individuals in each group is also very different: 1, 4, 7, 23, 23, 79. Thus, in this case it was not possible to assign hypothetical genotypes assuming 3 haplotypes. Again, we found no population structure underlying the six-way clustering. There was no correlation with latitude.

**Table S1:** Number of samples per inversion genotype

| Locus | Homop | Homoq | Heterozygous |
|---|---|---|---|
| 1 | 40 | 16 | 81 |
| 2 | 98 | 2 | 37 |
| 3 | 41 | 28 | 68 |
| 4 | 77 | 10 | 50 |
| 5 | 65 | 17 | 55 |
| 6 | 53 | 23 | 61 |
| 7 | 86 | 5 | 46 |
| 8 | 116 | 1 | 20 |
| 9 | 75 | 5 | 57 |

**Table S2:** Summary statistics from Fisher's Exact Test for $X^TX$ outliers inside and outside each locus region. Loci are arranged by descending *P*-value.

| Locus | estimate | conf.low | conf.high | p.value | A | B | C | D |
|---|---|---|---|---|---|---|---|---|
| 6 | 83.6 | 69.3 | 100 | $1.92\text{x}10^{-191}$ | 128 | 5372 | 2896 | 10158780 |
| 8 | 46.2 | 37.7 | 56.2 | $2.69\text{x}10^{-131}$ | 105 | 7895 | 2919 | 10156257 |
| 7 | 23.8 | 13 | 40.1 | $4.27\text{x}10^{-15}$ | 14 | 1986 | 3010 | 10162166 |
| 9 | 0 | 0 | 0.729 | 0.0122 | 0 | 17000 | 3024 | 10147152 |
| 4 | 0 | 0 | 0.918 | 0.0397 | 0 | 13500 | 3024 | 10150652 |
| 1 | 0 | 0 | 1.3 | 0.126 | 0 | 9500 | 3024 | 10154652 |
| 2 | 0.827 | 0.357 | 1.63 | 0.746 | 8 | 32492 | 3016 | 10131660 |
| 3 | 0 | 0 | 6.21 | 1 | 0 | 2000 | 3024 | 10162152 |
| 5 | 0 | 0 | 4.97 | 1 | 0 | 2500 | 3024 | 10161652 |

**Table S3:** Wilcox results $X^TX$; loci arranged by descending *P*-value.

| Chromosome | Loci | p value | w statistic | effect size |
|---|---|---|---|---|
| NW_022145609.1 | *Locus 8* | 8.59E-75 | 853333 | -0.5224987 |
| NW_022145603.1 | *Locus 6* | 2.62E-45 | 1440336 | 0.48466819 |
| NW_022145606.1 | *Locus 7* | 2.96E-10 | 224215 | -0.3291997 |
| NW_022145594.1 | *Locus 1, 2* | 3.57E-06 | 3047233 | 0.10383956 |
| NW_022145597.1 | *Locus 3* | 0.08141315 | 126353 | -0.1636074 |
| NW_022145600.1 | *Locus 4* | 0.21633735 | 984256 | 0.04884939 |
| NW_022145610.1 | *Locus 9* | 0.66112933 | 1224946 | 0.0142834 |
| NW_022145601.1 | *Locus 5* | 0.76384245 | 157523 | -0.0268429 |

**Table S4:** Chromosome name per Scaffold ID.

| Scaffold ID | Chromosome |
|---|---|
| NW_022145594.1 | 1 |
| NW_022145595.1 | 2 |
| NW_022145596.1 | 3 |
| NW_022145597.1 | 4 |
| NW_022145598.1 | 5 |
| NW_022145599.1 | 6 |
| NW_022145600.1 | 7 |
| NW_022145601.1 | 8 |
| NW_022145602.1 | 9 |
| NW_022145603.1 | 10 |
| NW_022145604.1 | 11 |
| NW_022145605.1 | 12 |
| NW_022145606.1 | 13 |
| NW_022145607.1 | 14 |
| NW_022145609.1 | 15 |
| NW_022145610.1 | 16 |
| NW_022145611.1 | 17 |
| NW_022145612.1 | 18 |
| NW_022145613.1 | 19 |
| NW_022145614.1 | 20 |
| NW_022145615.1 | 21 |

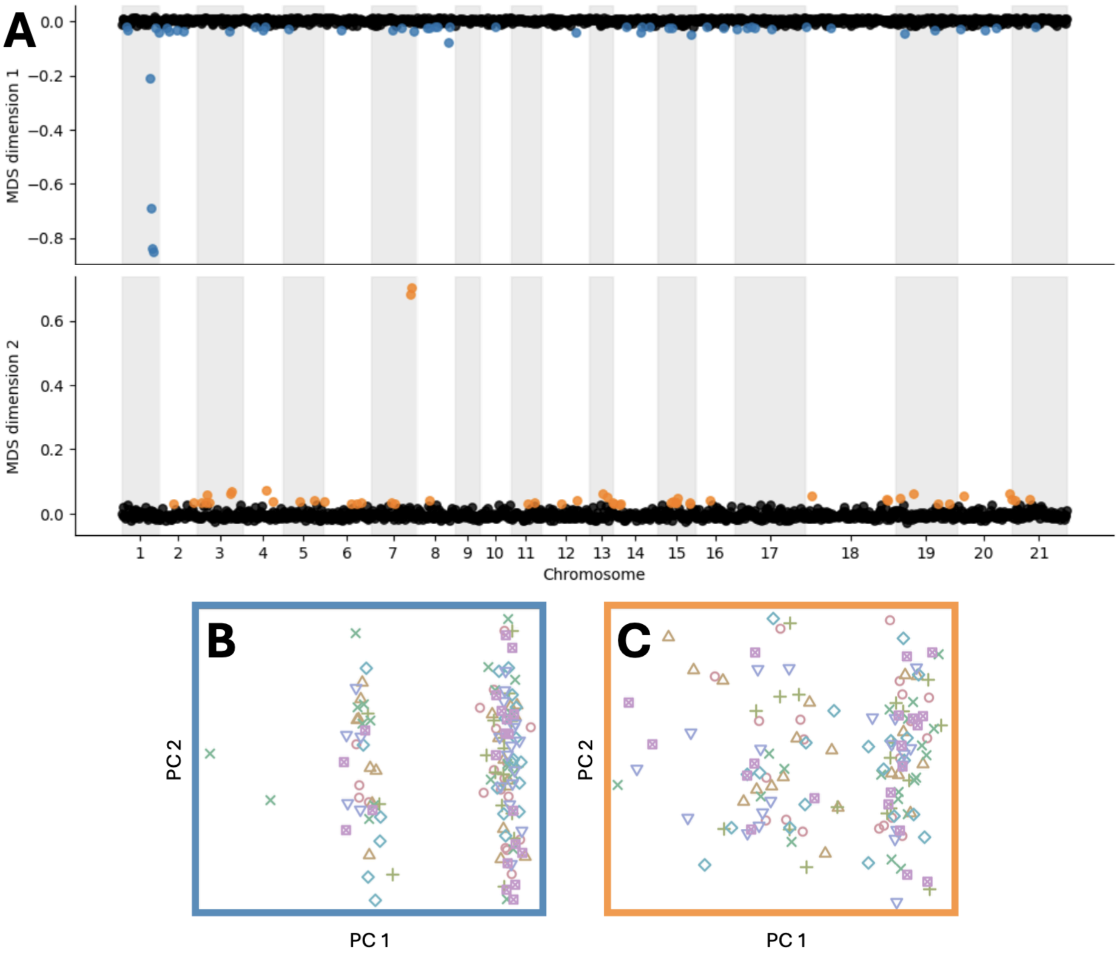


**Figure S1:** Local PCA across the whole genome, comparing genomic windows across chromosomes in a single MDS analysis. (A) shows the two MDS dimensions, with the 5\% outlier windows determined by the lostruct R package colored blue (for dimension 1) and orange (for dimension 2). (B) shows the PCA of the dimension 1 outlier windows, (C) shows the PCA of the dimension 2 outlier windows.

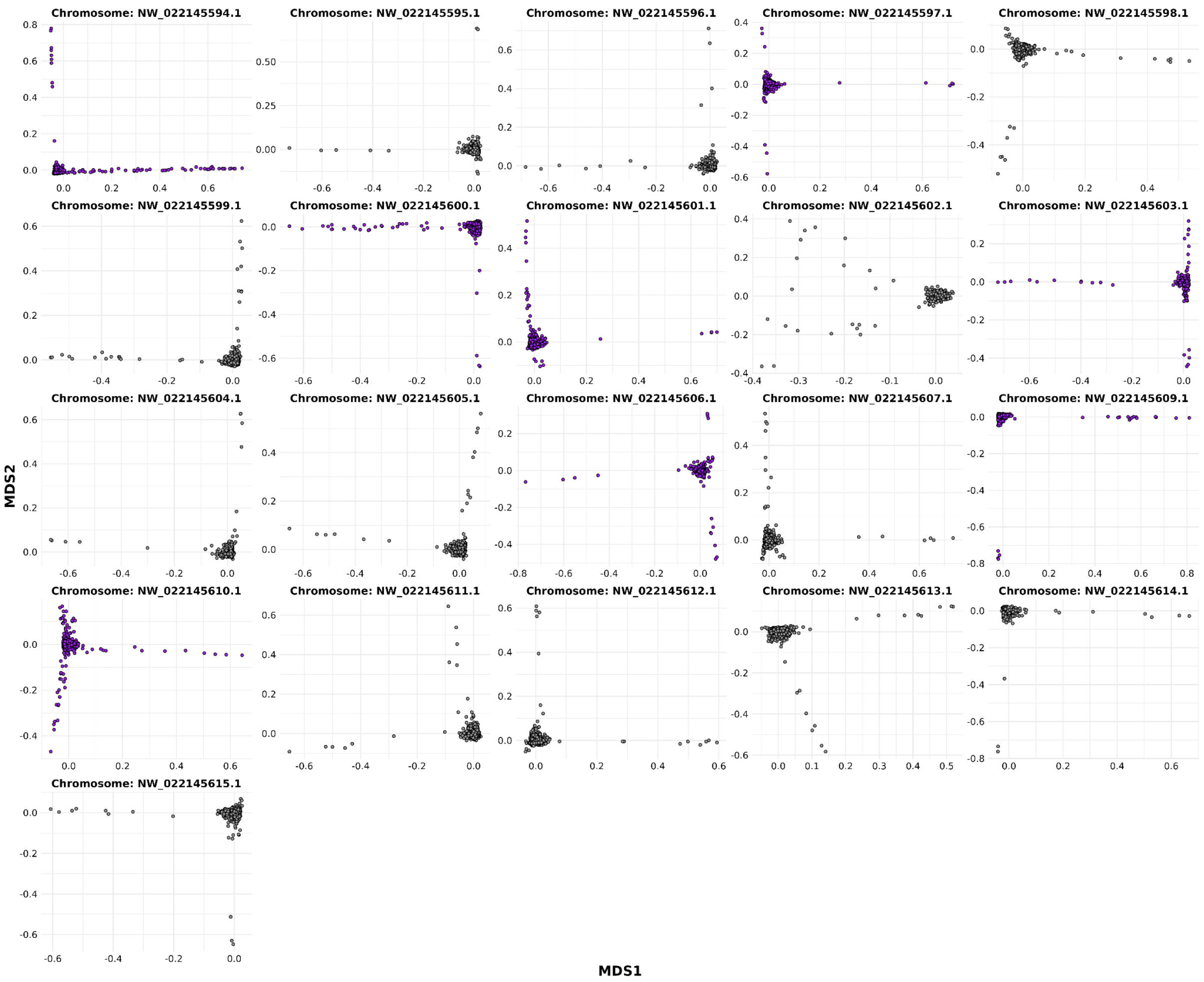


**Figure S2:** Multidimensional scaling (MDS) comparing windows on all chromosomes. Chromosomes with putative inversion polymorphisms based on the principal component analysis highlighted in purple. On each of the subplots the points (corresponding to windows) are arranged such that the biggest variation is along one of two dimensions. This means that the MDS is dominated by two large sources of variation and that outlier windows sharing similar PCs can be found by plotting each of the MDS dimensions along the genome.

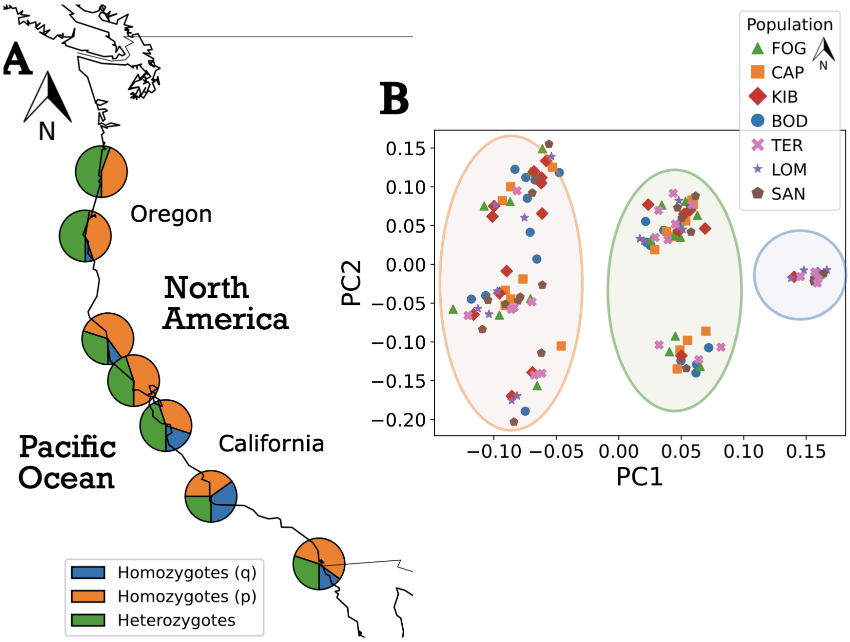


**Figure S3:** The frequency of individuals genotyped as heterokaryotypes at *locus 5* had a significant positive correlation with latitude ($R^2 = 0.64$, *P-value* = 0.0316), with more heterokaryotypes (green) in the North (A). PCA of genotype data spanning *locus 5* shows six distinct clusters, with no grouping by collection site.

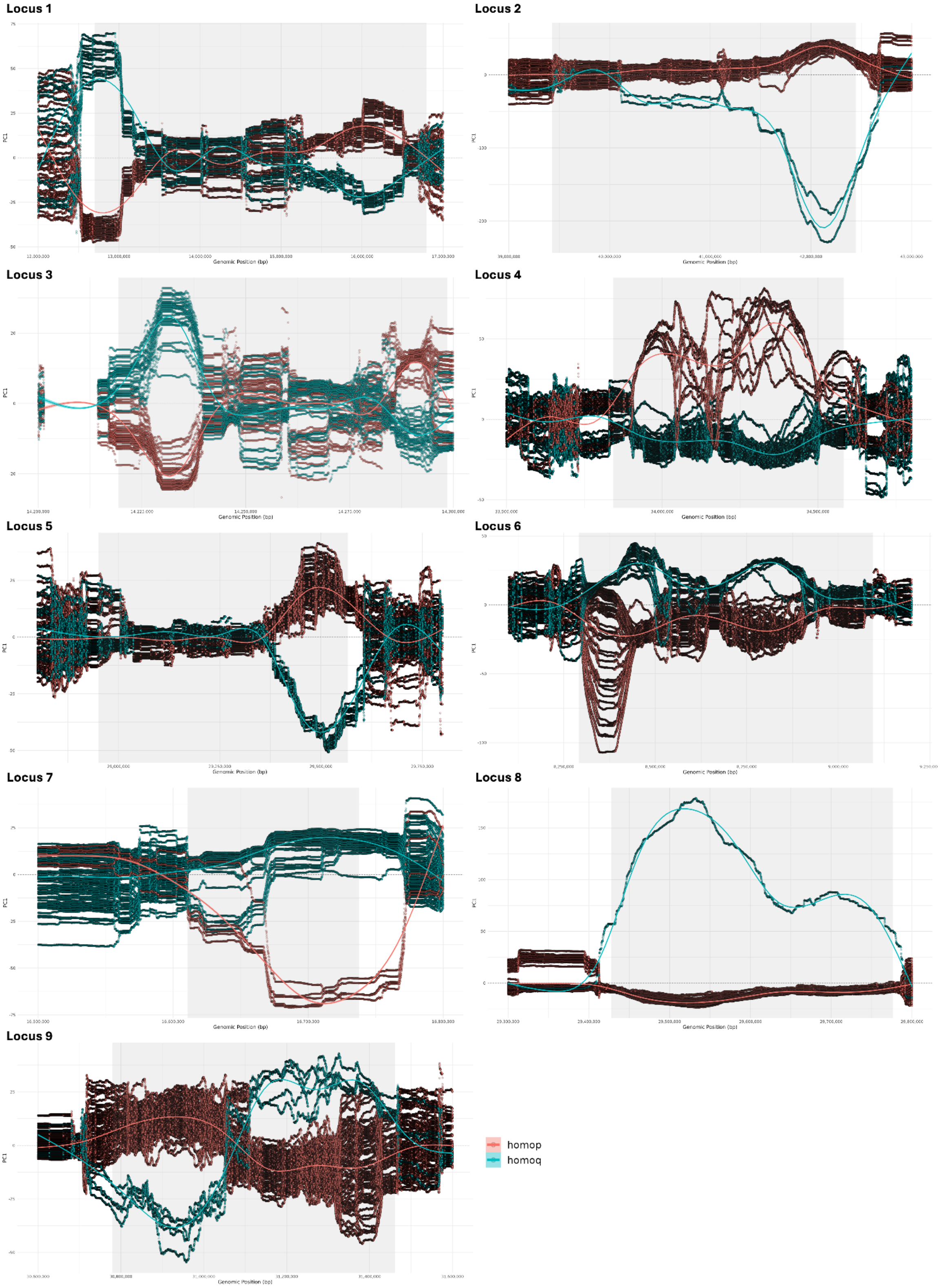


**Figure S4**: Chromosome wide windowed PCA using winPCA across loci of interest between homo-p and homo-q regions. Grey indicates the putative chromosomal inversion for each locus.

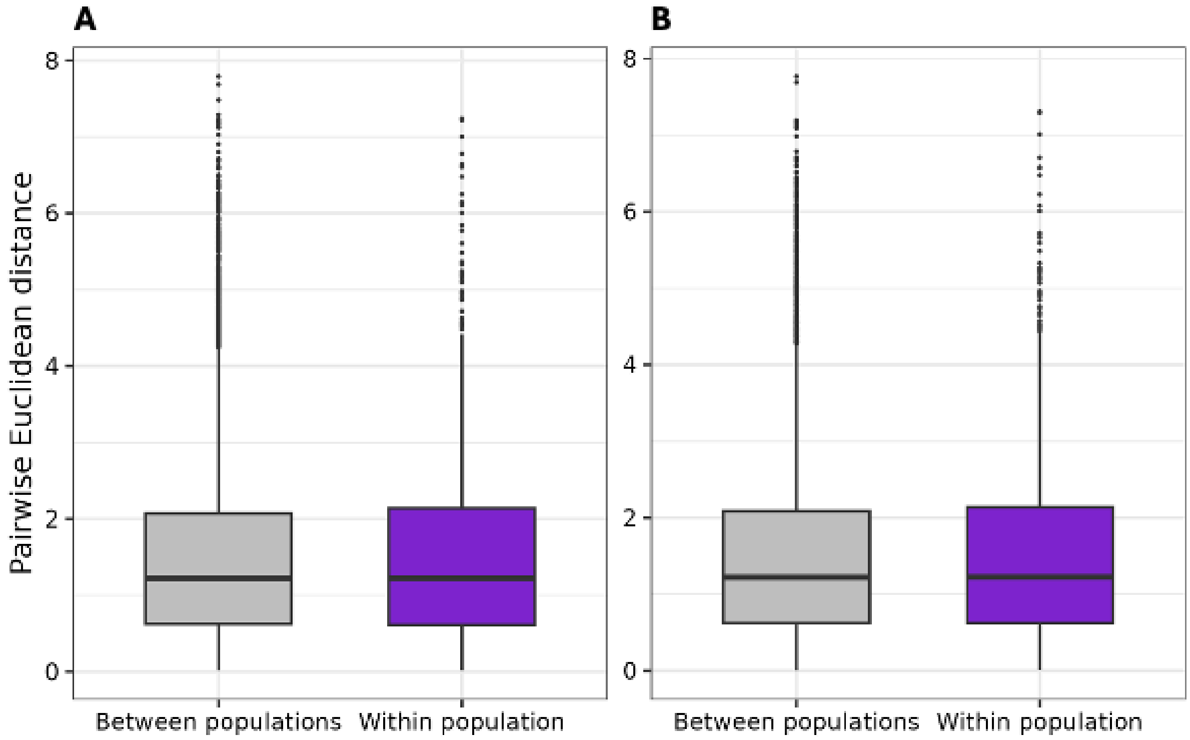


**Figure S5:** Comparisons of pairwise euclidean distance between populations and within populations for (a) all SNPs and (b) all SNPs minus those in putative inversions.

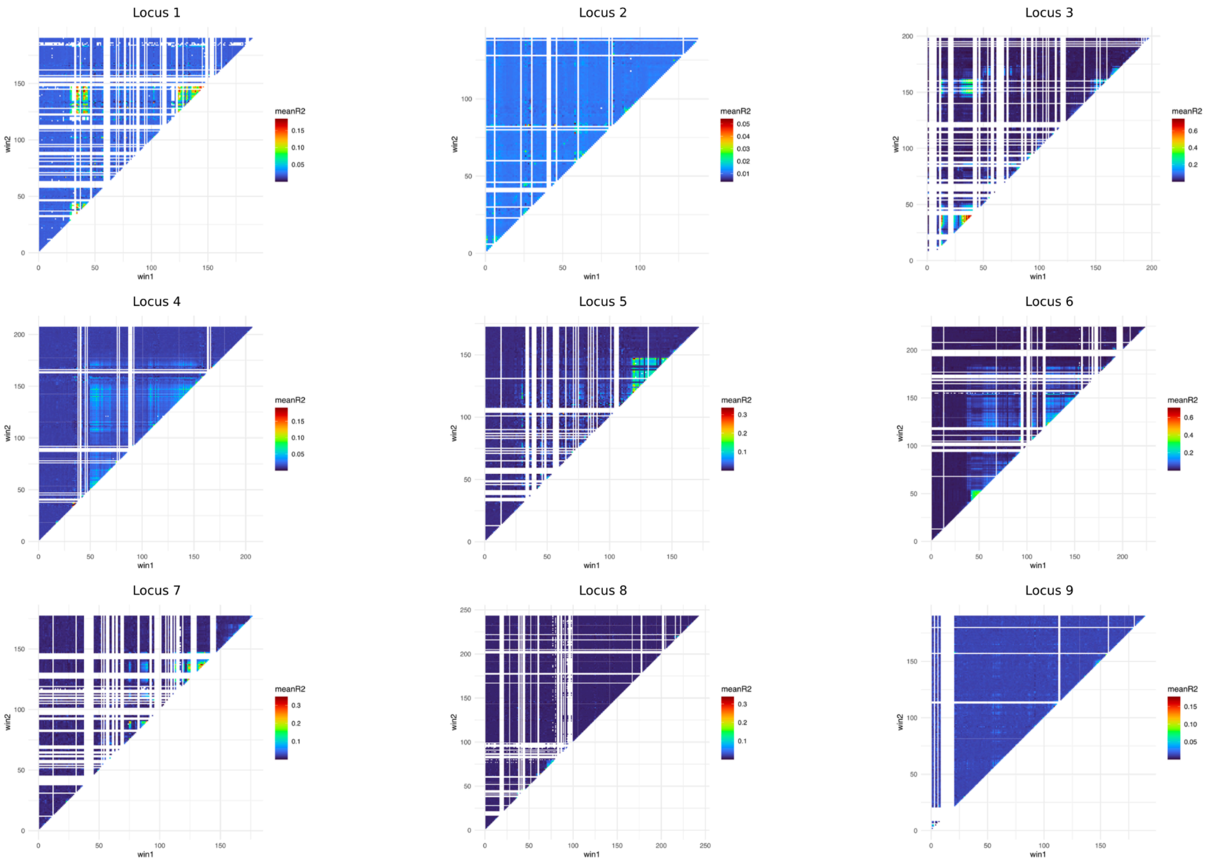


**Figure S6:** Linkage disequilibrium analysis for all nine loci when only including homokaryotypes (homop and homoq) in the analysis. Including only homop individuals resulted in significantly less linkage in all nine loci (data not shown), which is what we would expect for inversions recombining between chromosomes carrying the same inversion orientation.

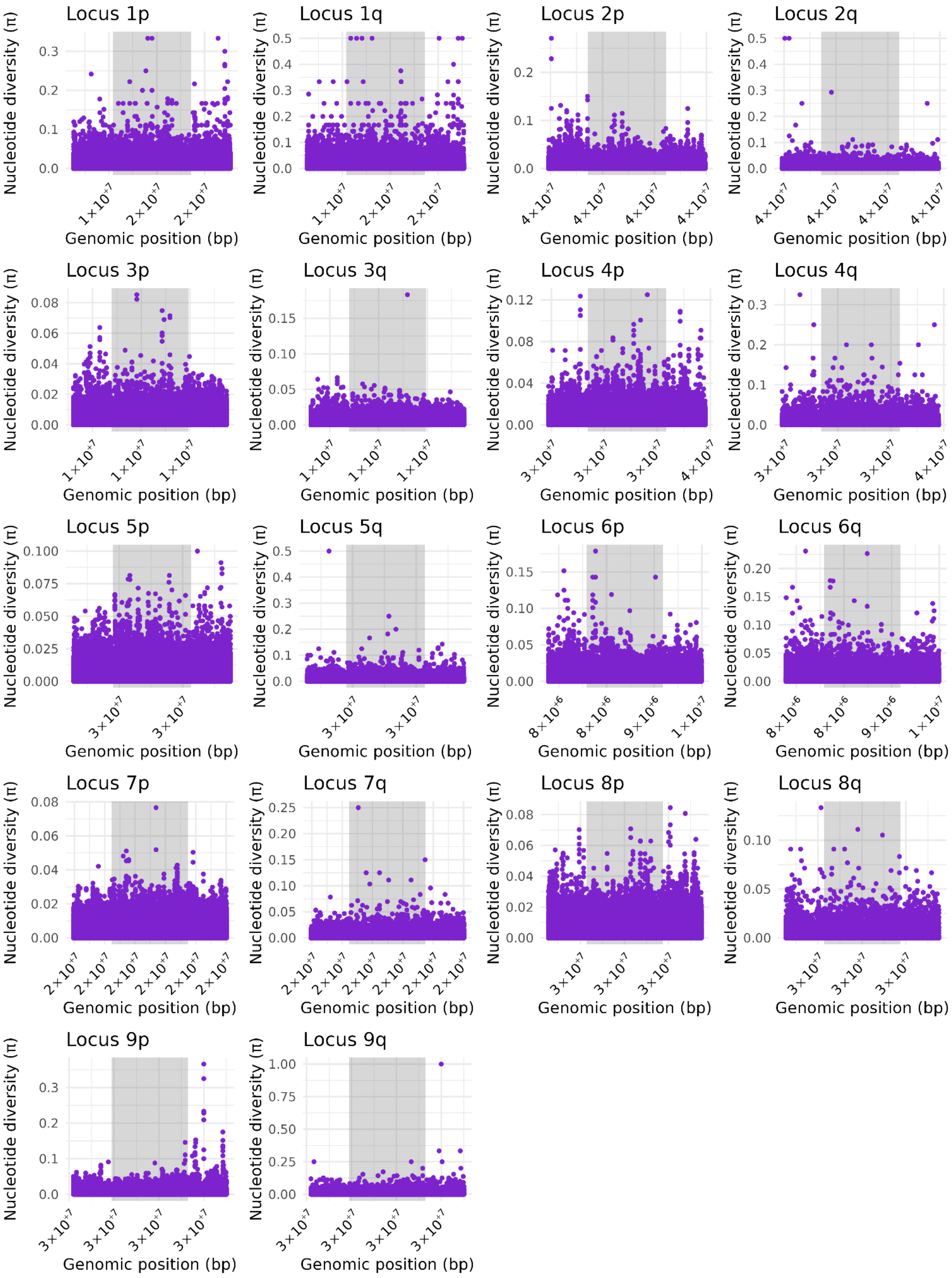


**Figure S7:** Nucleotide diversity (π) across putative inversions (*locus 1-9*). Loci are separated into two homokaryotype groups, *p* and *q*. Nucleotide diversity was calculated across 1000bp windows. Grey shading indicates putative inversion region.

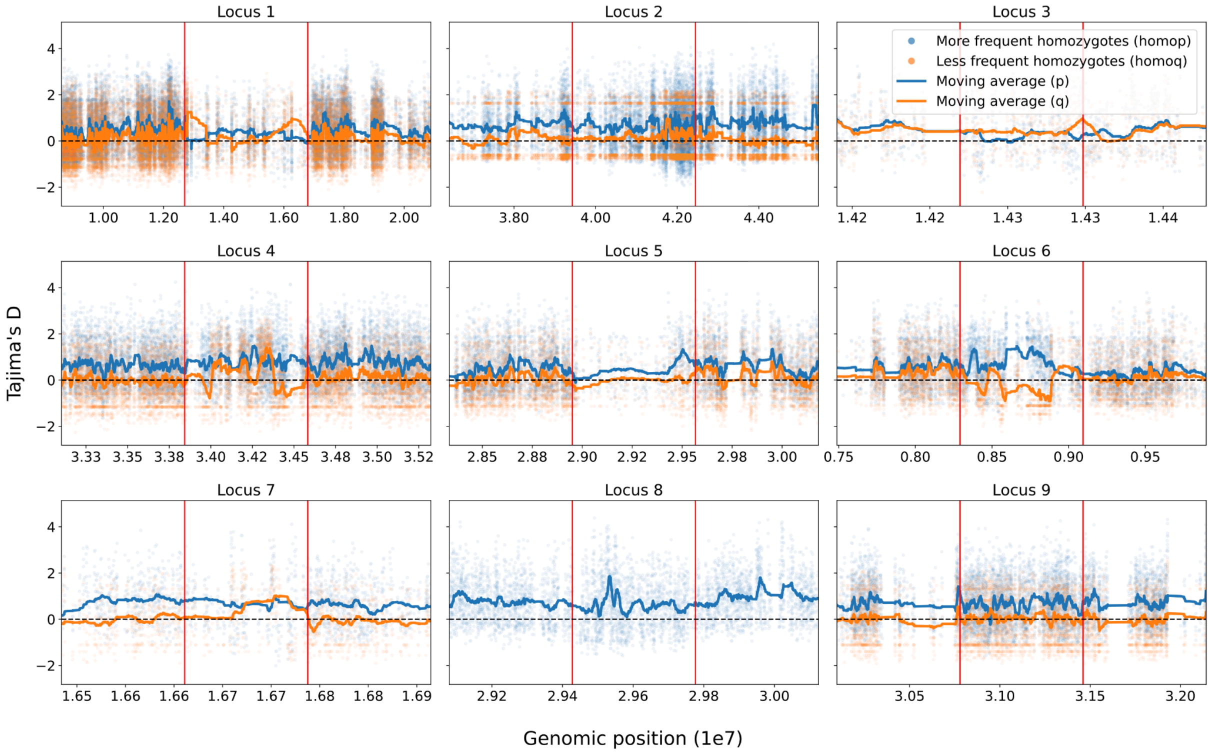


**Figure S8:** Tajima's D across each chromosome with a putative inversion polymorphism showing the two homokaryotype groups separately (blue and orange). Red lines mark the left and right putative inversion breakpoints. Semitransparent points show Tajima's D (window size = 100bp), solid lines on top show the moving average of those points. For inversion 8 there was only one individual in the less frequent homokaryotype group.

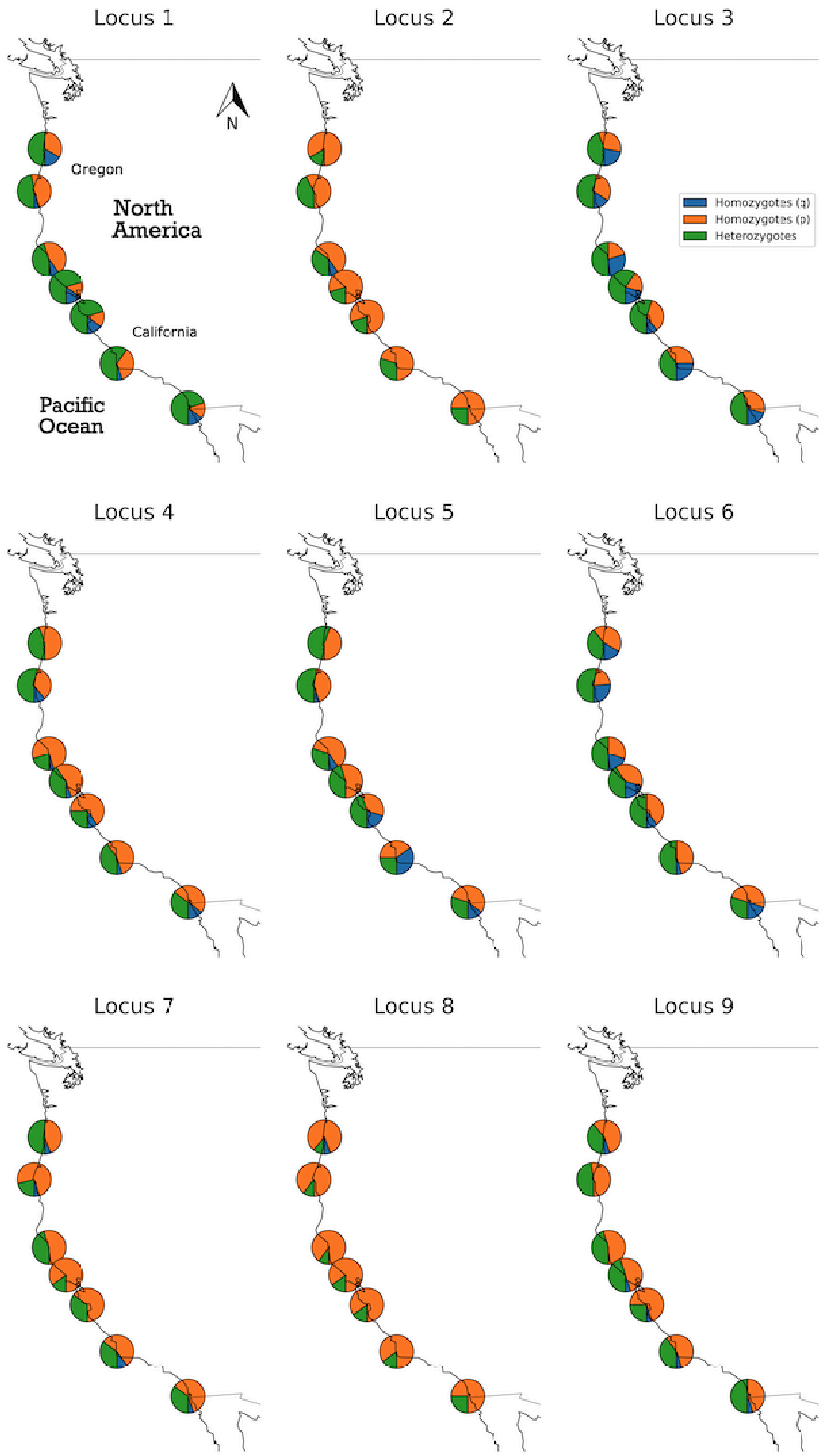


**Figure S9:** Frequency of homozygotes (blue and orange) and heterozygotes (green) for the nine putative inversions across collection sites. *Locus 1* and *5 were* correlated with latitude (*P-value* = 0.0015 *and P-value* = 0.0316, respectively).

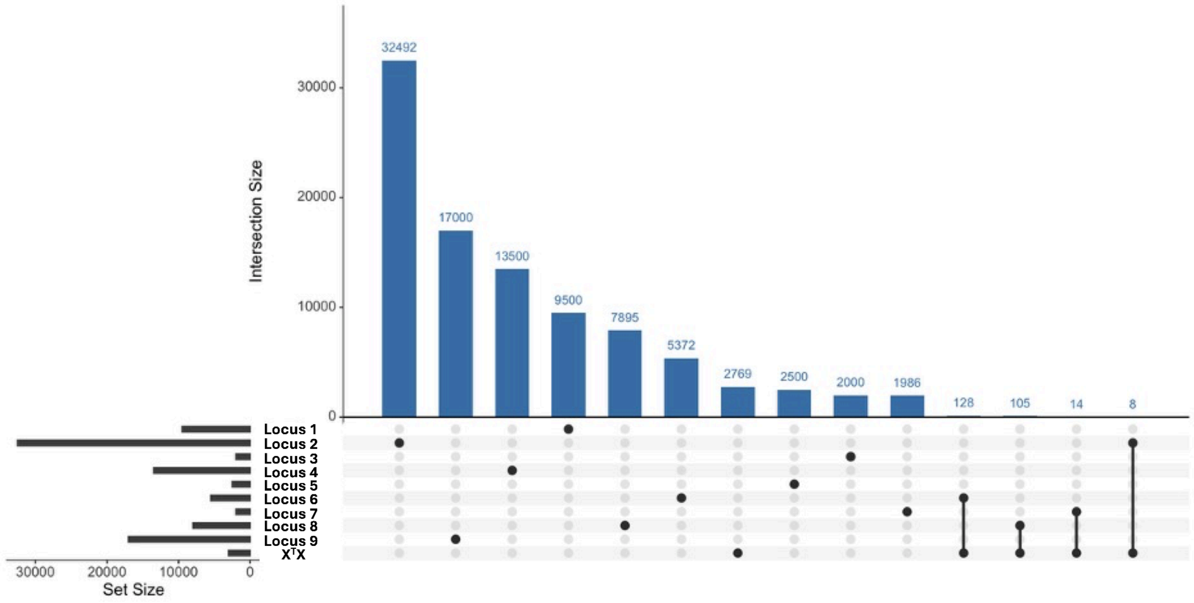


**Figure S10:** Upset plot of the number of SNPs shared across loci and those also identified as $X^TX$ outliers.
